# A Combined Human Gastruloid Model of Cardiogenesis and Neurogenesis

**DOI:** 10.1101/2022.02.25.481991

**Authors:** Zachary T. Olmsted, Janet L. Paluh

**Author notes:** Corresponding author: Dr. Janet L. Paluh, State University of New York Polytechnic Institute College of Nanoscale Science and Engineering, 4424 NanoFab East, 257 Fuller Road, Albany, NY 12203.

## Abstract

Multi-lineage development from gastruloids is enabling unprecedented opportunities to model and study human embryonic processes and is expected to accelerate *ex vivo* strategies in organ development. Reproducing human cardiogenesis with neurogenesis in a multi-lineage context remains challenging, requiring spatiotemporal input of paracrine and mechanical cues. Here we extend elongating multi-lineage organized (EMLO) gastruloids to include cardiogenesis (EMLOC) and describe interconnected neuro-cardiac lineages in a single gastruloid model. Contractile EMLOCs recapitulate numerous interlinked developmental features including heart tube formation and specialization, cardiomyocyte differentiation and remodeling phases, epicardium, ventricular wall morphogenesis, chamber-like structures and formation of a putative outflow tract. The EMLOC cardiac region, which originates anterior to gut tube primordium, is progressively populated by neurons in a spatial pattern mirroring the known distribution of neurons in the innervated human heart. This human EMLOC model represents the first multi-lineage advancement for the study of coincident neurogenesis and cardiogenesis.

## INTRODUCTION

The complex nature of *in vivo* cardiogenesis underlies the difficulties in establishing *in vitro* cardiac developmental models with human cells. The heart is the first organ to form in the mammalian embryo, caudal to the embryonic brain and within the developing trunk. It becomes contractile as a tube prior to complex morphogenesis into septated chambers and co-developmental population by neurons for innervation (Harvey, 2002; Hasan, 2013). In order to accommodate both contractility and structural rearrangement, the developing heart undergoes alternating phases of cardiac differentiation and morphogenesis (Ivanovitch et al., 2017). Calcium handling properties become refined during cardiac differentiation (Tyser et al., 2016). The cardiac crescent is the first bilateral structure to form and precedes epithelialization and formation of the transversal heart tube. At this stage, the heart tube remains open at the dorsal aspect, bound by dorsal mesocardium, and then seals during formation of the closed linear heart tube and outflow tracts. Intrinsic cell-driven forces within the tube and extrinsic physical constraints are known to mediate the establishment of left-right asymmetries required for heart function (Desgrange et al., 2018). Such complexity in cardiogenesis lays the framework for lifelong functioning of the adult heart, but also underlies the propensity for congenital heart disease in humans where developmental errors induce cardiac malformations (van der Linde et al., 2011; Desgrange et al., 2018). The ability to generate *in vitro* models of heart development that mimic essential aspects of multi-lineage input to cardiogenesis will benefit biomedical treatments of heart disease and progress towards *ex vivo* organogenesis.

Organoid technology is revolutionizing the study of human development and disease, recapitulating key aspects of spatiotemporal tissue morphogenesis (Clevers, 2016; Olmsted and Paluh, 2021c). Most current organoid technologies are directed towards single tissue endpoints that lack the cellular contextual diversity present in normal organogenesis through inductive and mechanical interactions. As such, the ability to generate organotypic human cardiac organoids that form according to the *in situ* developmental signaling blueprint and integrate with the developing nervous system has not been achieved. The existing human cardiac organoid models derive primarily from pre-differentiated cardiomyocytes and their spheroid aggregates that form irrespective of developmental timelines (Nguyen et al., 2014; Giacomelli et al., 2017; Polonchuk et al., 2017; Andersen et al., 2018), or models that rely on integrated bioengineering efforts to constrain morphogenetic patterning (Ma et al., 2015; Lind et al., 2017; Macqueen et al., 2018; Hookway et al., 2019). These models lack identified critical inductive tissues indispensable to natural heart development such as the foregut, described as a central organizer of cardiogenesis in multiple species and acting through both inductive and structural interactions between endoderm and splanchnic mesoderm (Nascone and Mercola, 1995; Schultheiss et al., 1995; Varner and Taber, 2012; Anderson et al., 2016; Kidokoro et al., 2018; Han et al., 2020).

Two recent studies with human iPSCs succeeded in the co-production of cardiac and gastrointestinal tissue in single organoids without organized chambers (Silva et al., 2021; Drakhlis et al., 2021). As well, Hofbauer et al. (2021) succeeded in generating self-organized, isolated cardioids exhibiting chamber-like structures from human pluripotent stem cells that were used to model cardiac injury (Hofbauer et al., 2021). Although important advances to the cardiogenesis field, neural cells were not co-generated in these systems and were absent. One murine study generated chambered cardiac organoids from mESCs by embedding in exogenous extracellular matrix (ECM) with supplied FGF4 (Lee et al., 2020). More recently, Rossi et al. (2021) used mESC-derived gastruloids to recapitulate aspects of early cardiogenesis including first and second heart field contributions without extracellular matrix (ECM) embedding. Gastruloid research has been broadly applicable for conducting multi-lineage interaction studies in the trunk (van den Brink et al., 2014; van den Brink et al., 2020; Veenvliet et al., 2020; Olmsted and Paluh, 2021a). However, no study with human cells has succeeded in generating a *de novo* model to recapitulate cardiogenesis in an embryo-like, multi-lineage context and, in particular, with neuronal cooperative development that is a vital functional component. Here we achieve this goal and describe human gastruloids that capture numerous key developmental aspects of human cardiogenesis and neurogenesis along with endoderm-derived primitive gut tube and other lineages.

We recently described a unique human trunk model system referred to as elongating multi-lineage organized (EMLO) gastruloids (Olmsted and Paluh, 2021a, 2021b). Neural crest lineage in EMLOs reveals insights into enteric development and the formation of the enteric nervous system. The enteric multi-lineage niche in EMLOs achieved only limited embryonic cardiogenesis that included generation of cardiomyocytes anterior to the gut tube. We therefore hypothesized that EMLOs could be coaxed developmentally, if provided the necessary cues, towards more extended cardiac differentiation with reproducible morphogenesis. We reoptimized our EMLO formation protocol to include angiocrine and pro-cardiogenic factors previously detailed (Rossi et al., 2021). We now demonstrate a gastruloid strategy for neuro-cardiac co-developed tissues that recapitulate aspects of early human heart morphogenesis with neuronal integration. We track multiple events in cardiomyocyte differentiation from splanchnic mesoderm and observe spontaneous contractility, chamber precursor formation, early constrictions and septations, epicardium, and putative structures resembling the outflow tracts. Critically, EMLOCs not only retain the interacting neural compartment but achieve neurogenesis to generate an organized co-developed neuro-cardiac gastruloid. This work establishes EMLOCs as an advanced model for human cardiogenesis and the integration with endoderm and neurons towards the goal of organ innervation.

## RESULTS

### Angiocrine and pro-cardiogenic factors redirect multi-lineage EMLO gastruloids for optimized human developmental cardiogenesis (EMLOCs)

We previously generated EMLO gastruloids with co-developing central and peripheral neurons and trunk mesendoderm including components of the enteric nervous system (Olmsted and Paluh, 2021a, 2021b). To test the ability of EMLOs to model human developmental cardiogenesis (EMLOCs), we modified exposure to growth factors during early formation and polarization stages in a revised protocol (**Figure 1A, Figure S1**). EMLOCs were generated with the hiPSC line H3.1.1 that we previously validated for differentiation into contractile cardiomyocytes (Tomov et al., 2016) and that forms EMLOs by the original protocol described (Olmsted and Paluh, 2021a, 2021b). EMLOCs were handled identically to EMLOs during a 2D induction phase up to 48 h post-aggregation in shaking cultures. At 48 h, the N2B27 supplemented with 10 ng/ml FGF2, 2 ng/ml HGF, and 2 ng/ml IGF-1 in EMLOs was instead supplemented with 30 ng/ml FGF2, 5 ng/ml VEGF, and 0.5 mM ascorbic acid (AA) (**Figure S1A-C**) as was done for the mESC cardiac gastruloid model (Rossi et al., 2021). The new cardiogenic factors were added in the absence of the initial factors used for induction. The EMLOC gastruloids were maintained in this pro-cardiogenic medium to day 7. In the first 48 h prior to the medium change, early germ layer biomarkers were expressed in appropriately sized gastruloids as expected for the EMLO protocol. That is, uniform expression of SOX2 24 h post-aggregation, with little to no expression of GATA6 and heterogenous expression of FOXA2 (**Figure S1D**). Mitotically active cells in the gastruloid were visible (**Figure S1E**). After the medium change, the EMLOC gastruloids that were continually maintained in shaking culture began to elongate by day 5. By day 7, elongated gastruloids formed thin-walled, dilated chamber-like structures with spontaneous contractility (**Figure 1; Movies S1 and S2**). Partitioning of cardiogenic chamber precursors was visible by phase contrast microscopy (**Figure S1F**), and validated by expression of the early cardiogenic transcription factor GATA6 along with cardiac Troponin T (cTnT) (**Figure S1G**). Multiple distinct cavities resembling chambers were also identified by immunofluorescence in single EMLOCs with early evidence of septation (**Figures 1B and 1C**). These findings support the ability to direct cardiogenic tissue precursors within gastruloids by early manipulation of growth conditions and signaling factors, demonstrated by the genesis of self-organizing cardiogenic compartments. We next performed single cell RNA sequencing (scRNAseq) to further delineate the cell and tissue precursor types generated by this protocol.

**Figure 1.**
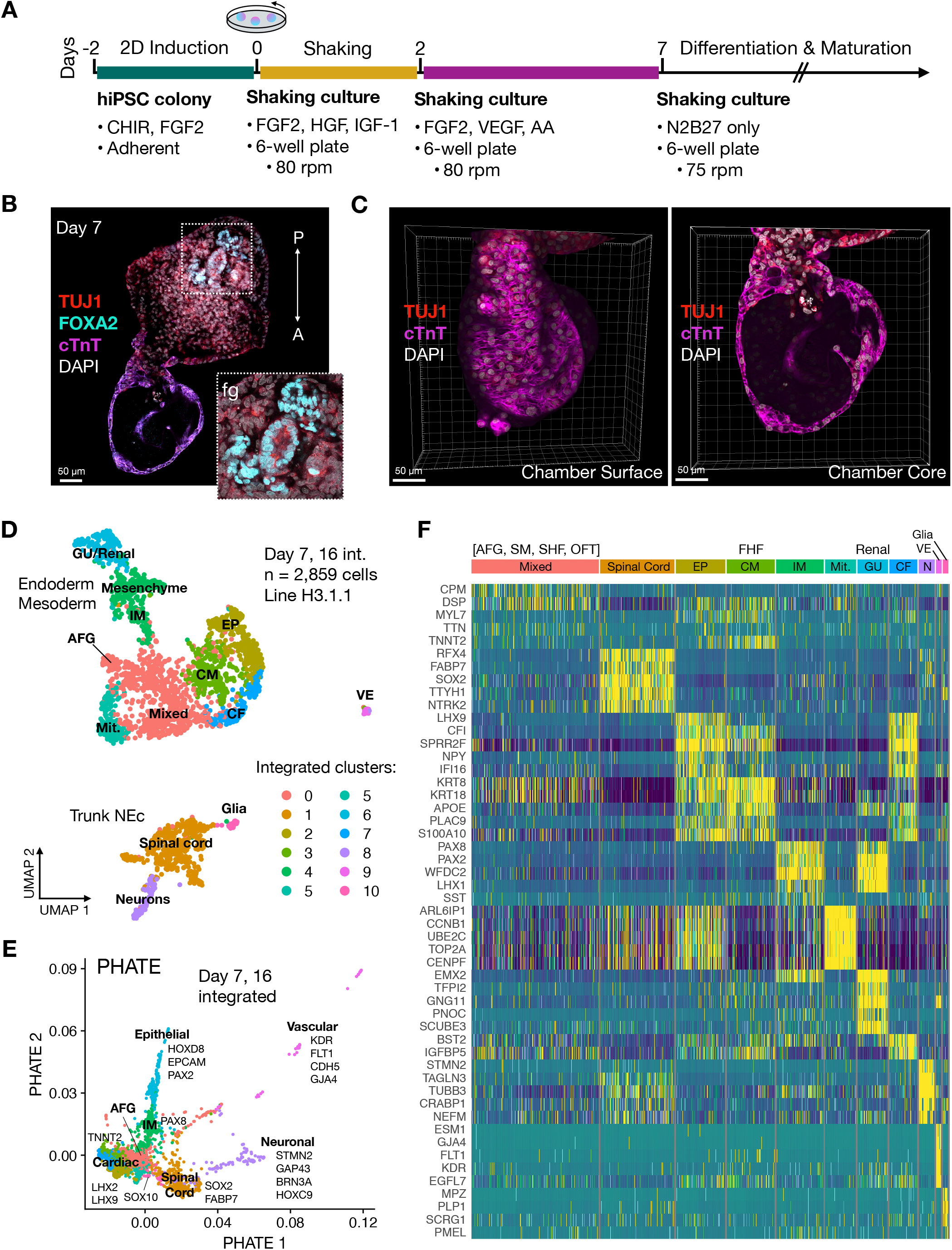
EMLOC gastruloids capture cell type diversity of human cardiogenesis and trunk development. (**A**) Overview of protocol for EMLOC gastruloid generation. Cardiogenesis was induced at 48 h post-aggregation by addition of VEGF and ascorbic acid (AA). (**B**) Immunofluorescence of day 7 H3.1.1 EMLOC immunostained for TUJ1 (red), cTnT (magenta), FOXA2 (cyan) and counterstained with DAPI (grey). Inset is high magnification Z-slice of FOXA2 foregut (fg) initialization. Anterior (A) to posterior (P) orientation is descriptive. (**C**) 3D reconstruction of anterior cardiac region from (**B**) with TUJ1 (red), cTnT (magenta) and DAPI (grey). The EMLOC chamber surface (left) and core (right) are shown. Individual scale bars provided. (**D**) UMAP visualization of ten annotated clusters from the integrated scRNAseq dataset containing day 7 (1,004 cells) and day 16 (1,855 cells) time points in EMLOC formation (2,859 total cells). (**E**) PHATE visualization from the integrated dataset shown in (**D**). Cell lineages are labeled along with upregulated DEGs. (**F**) Heatmap of the top five DEGs for each cluster of the integrated scRNAseq dataset (see also **Table S1**, **Supplementary data file**). Bright yellow depicts upregulated differential gene expression. Abbreviations: anterior foregut (AFG), cardiac fibroblast (CF), cardiomyocyte (CM), epicardial cells (EP), first heart field (FHF), genitourinary (GU), intermediate mesoderm (IM), mitotic (mit), neuronal (N), outflow tract (OFT), second heart field (SHF), splanchnic mesoderm (SM), vascular endothelial cells (VE) .

### EMLOC gastruloids generate diverse embryonic cell types of the human trunk revealed by scRNAseq analysis

Single cell sequencing of H3.1.1 derived EMLOCs was performed at two time points that are day 7 and day 16 after initial aggregation in shaking culture. The integrated dataset of both time points was analyzed (2,859 cells) (**Figures 1D-1F**) along with each time point individually (day 7: 1,004 cells; day 16: 1,855 cells) (**Figure 2, Figure S2**). The integrated dataset was generated in Seurat and visualized using UMAP and PHATE methods. Ten clusters were produced and were annotated using differentially expressed genes (DEGs) and cell or tissue type characteristic biomarkers from the literature (**Figures 1D and 1E)**. The top five DEGs for each cluster of the integrated dataset is provided (**Figure 1F**) in addition to a comprehensive list of DEGs and genes used for cluster annotation (**Table S1**; **Supplementary data file**). Clusters were annotated as trunk neuroectoderm/spinal cord progenitors (cluster 1), trunk neurons (cluster 8), peripheral glia/Schwann cells (cluster 10), mitotic cells (cluster 5), mixed cell types (cluster 0), cardiomyocytes (CM; cluster 3), epicardial cells and cardiac fibroblasts (EP, CF; clusters 2 and 7), intermediate mesoderm and metanephric mesenchyme (IM; cluster 4), genitourinary/renal epithelium (GU; cluster 6), and vascular endothelium (VE; cluster 9). By PHATE analysis of day 16 cells, we visualized distinct cardiac, epithelial and neural lineages (**Figure 2A**). We overlaid important signaling pathways including BMP, SHH, and WNT signaling that were delineated along distinct lineages (**Figure 2B**). *BMP4* was highly expressed in the cardiac region while *BMP7* bifurcated along neural and trunk epithelial lineages. *SHH* was upregulated in a region within cluster 0 of anterior foregut (AFG) phenotype (*FOXA2*, *NKX2-1*, *SHH*, *EPCAM*) that is a known developmental organizer of cardiogenesis (Anderson et al., 2016). *WNT2B* expression localized to the cardiac region while *WNT1* and *WNT3A* localized to the spinal cord region with known involvement in neural tube morphogenesis and neural crest patterning. *WNT1* is shown. Cadherin and *HOX* genes were also delineated along respective lineages, consistent with developmental cadherin and *HOX* codes *in vivo* (**Figure 2C**). *CDH11* was upregulated in the cardiac region while *CDH6* was upregulated in neuroectoderm and mesenchyme and *CDH1* was upregulated in epithelium. Distinct expression of *HOXC6* and *HOXC9* in neural clusters was indicative of caudal neuraxis and trunk spinal cord. *HOXD8* and *HOXD9* were specific to epithelium with renal mRNA expression phenotype (cluster 6) while *HOXA4* was predominantly expressed in the cardiac region. This scRNAseq analysis reflects a diversity of cell and tissue precursor types generated within EMLOCs with signaling networks, adhesion proteins, and transcription factors mirroring *in vivo* development. We elaborate on specific features of the annotated clusters throughout the manuscript where indicated, with emphasis on neural, cardiac, and foregut endodermal lineages.

**Figure 2.**
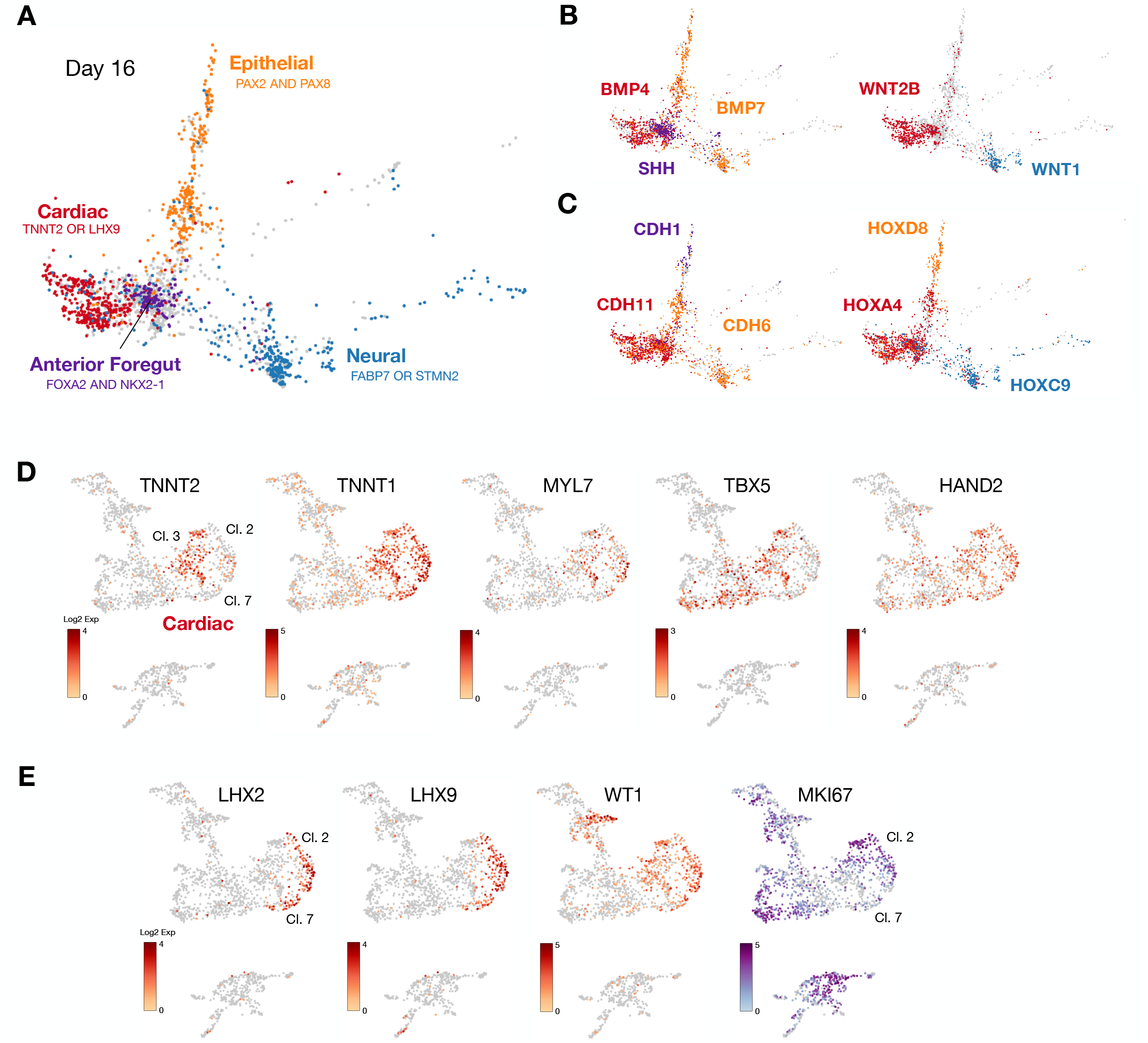
Diverging lineages in EMLOCs advance according to *in vivo* developmental principles. (**A**) Day 16 EMLOC scRNAseq dataset visualized by PHATE. Superimposed developmental lineages include cardiac (*TNNT2* or *LHX9*), anterior foregut (*FOXA2* and *NKX2-1*), epithelial (*PAX2* and *PAX8*), and neural (*FABP7* or *STMN2*). (**B**) Signaling pathways segregate along diverging lineages in EMLOCs. *BMP4* vs. *BMP7* vs. *SHH* is shown (left) along with *WNT2B* vs. *WNT1*. (**C**) Preserved cadherin and Hox codes in EMLOCs. *CDH1* vs. *CDH6* vs. *CDH11* is shown (left) along with *HOXA4* vs. *HOXC9* vs. *HOXD8* (right). (**D-E**) Day 16 EMLOC scRNAseq dataset visualized by UMAP. (**D**) Genes for sarcomere proteins involved in contractility (*TNNT2, TNNT1, MYL7*) are upregulated primarily in cluster 3. Biomarkers for FHF (*TBX5*) and SHF (*HAND2*) are shown. (**E**) Epicardial cell and cardiac fibroblast biomarkers *LHX2*, *LHX9*, *WT1* are upregulated in clusters 2 and 7. Cell proliferation marker *MKI67* depicts fewer mitotic cells in cluster 7 vs. cluster 2.

### Multiple derivatives of splanchnic mesoderm in EMLOCs identified by scRNAseq

Cardiogenic mesoderm gives rise not only to working cardiomyocytes but also contributes to epicardium, endocardium, connective tissue, outflow tract, valves and the conduction apparatus. Cluster annotation and analysis of scRNAseq data in day 7 and day 16 EMLOCs identified diverse cell types involved in cardiogenesis arising from splanchnic mesoderm (**Figures 2D-2E, Figure S3**) including cardiomyocytes that form myocardium, cells of the proepicardium and epicardium, and cardiac fibroblasts, in addition to endocardium and vascular endothelium. In mouse, cardiac precursors arise within the splanchnic mesoderm and differentiate into cardiomyocytes by assembling contractile machinery (Tyser et al., 2016). EMLO gastruloids were previously shown to contain splanchnic mesoderm permissive to cardiomyocyte differentiation (Olmsted and Paluh, 2021a). Similarly, EMLOCs retain characteristic expression of the splanchnic mesoderm biomarkers *FOXF1*, *PDGFRA*, *TWIST1*, and *PRRX2* (**Figure S3A**). *GATA4* and *GATA6* that are indispensable for cardiogenesis were expressed in a similar distribution (**Figure S3B**). Sarcomeric proteins associated with cardiomyocytes such as *TNNT2* (cardiac troponin), *TNNT1* (slow type troponin) and *MYL7* were most localized to cluster 3 at day 16 along with first heart field (FHF) and second heart field (SHF) biomarkers *TBX5* and *HAND2* (**Figure 2D**). Additional FHF genes (*NKX2-5*, *HAND1*) and SHF genes (*MEF2C*, *ISL1*, *TBX18*) were similarly distributed. At the day 16 time point, cardiomyocytes were primarily of ventricular cell phenotype. DEGs involved in regulating ventricle growth, morphogenesis, and contractility were identified (**Figure S3B**). Cluster 3 contained additional upregulated genes involved in fetal heart development such as dimeric *KRT8*/*KRT18*, *APOE*, *PLAC9* and *S100A10*. These genes were also upregulated in cluster 2 that were annotated as epicardial (EP) cells based on expression profile (*WT1*, *TCF21*, *TPJ1*, *LHX2*, *LHX9*, *TBX18, PLAC9*) (**Figure 2E**). Cluster 7 had a similar expression profile to cluster 2, but with several distinguishing features such as reduced proliferation assessed by *MKI67*, upregulation of *IGFBP5* associated with cardiac fibroblast activation and *BTS2* that interacts with *IFI16*, a biomarker of mature cardiac fibroblasts. A previous analysis of human fetal cardiogenesis revealed subclusters of actively proliferating, early fibroblasts and those enriched for ECM organization with less proliferation (Cui et al., 2019). A similar observation was made here between cluster 2 (high proliferation by *MKI67*) and cluster 7 (low proliferation by *MKI67*). The scRNAseq clusters representing cardiogenesis expressed a characteristic milieu of cardiomyocyte and fibroblast-derived ECM genes (**Figure S4**). Cluster 0 contained upregulated genes indicating a mixture of cell types from cardiogenic mesoderm including contractile cells (*TNNT2*, *MYL7*), outflow tract cells (*PDE5A*, *ISL1*, *FN1*, *MEGF6, MSX2, SEMA3C, EMILIN1, CNN1, TAGLN*), and cells involved in atrioventricular conduction and organization (*GJA1*, *CACNA1H, TBX3*, *CXCL12*, *DSP*). Anterior foregut progenitor cells were also identified (*HHEX, SHH, FOXA2*). Cells expressing cardiac neural crest biomarkers (*ETS1, EDNRA, TGIF1, HOXA3*) were dispersed throughout the four clusters (**Figure S3C**). Cardiac neural crest cells *in vivo* play critical roles in cardiogenesis and organization, including valve and outflow tract contributions. Regulatory subnetworks within EMLOCs therefore generate a range of the cell types involved in cardiogenesis along with an appropriate ECM milieu.

### Human EMLOC multi-lineage gastruloids form chamber-like structures with spontaneous contractility and calcium signaling

Intracellular changes in Ca^2+^ couple cardiomyocyte depolarization with contraction. To demonstrate that EMLOCs express calcium-regulated contractile proteins and achieve calcium-mediated contractility, we performed 3D fixed and live cell imaging analysis (**Figure 3**). The cardiogenic compartment in EMLOCs exhibited visible sarcomeres using cTnT immunofluorescence (**Figure 3A**) and spontaneous contractility (**Figure 3B**). To quantify spontaneous contractility, we compared beating phenotypes in EMLOC-directed gastruloids versus the original EMLO protocol that is not optimized for cardiogenesis (Olmsted and Paluh, 2020a) (N = 3 repeat experiments; ***p = 0.0005, t = 10.40 df = 4 by unpaired two-tailed t-test). We further performed live calcium imaging with Fluo-4 AM (**Figures 3C-3E**). In **Figure 3C**, we demonstrate robust Fluo-4 AM activity in the contractile region of two gastruloids and quantify F/F_o_ calcium transients (**Figure 3D; Movies S3 and S4**). The EMLOCs shown were captured in the same field and are representative of the quantified population. Median calcium transients per min (corresponding to beat frequency) was 10.5 (n = 10, max = 19, min = 5, median = 10.5, q1 = 8, q3 = 12.75) (**Figure 3E**). Known genes involved in cardiac action potential conduction and calcium handling are expressed, including *IRX3* and *IRX5* that play roles in rapid ventricular conduction and cardiac repolarization, respectively, along with the *ITPR2* calcium channel and sodium-calcium exchanger *SLC8A1*/NAC1 that were all expressed with similar distribution (**Figure 3F**). Together these data demonstrate that morphological cardiogenesis chamber features, calcium signaling and spontaneous contractility can be directed in human EMLOCs by modifying the EMLO protocol with exposure to angiocrine and pro-cardiogenic growth factors.

**Figure 3.**
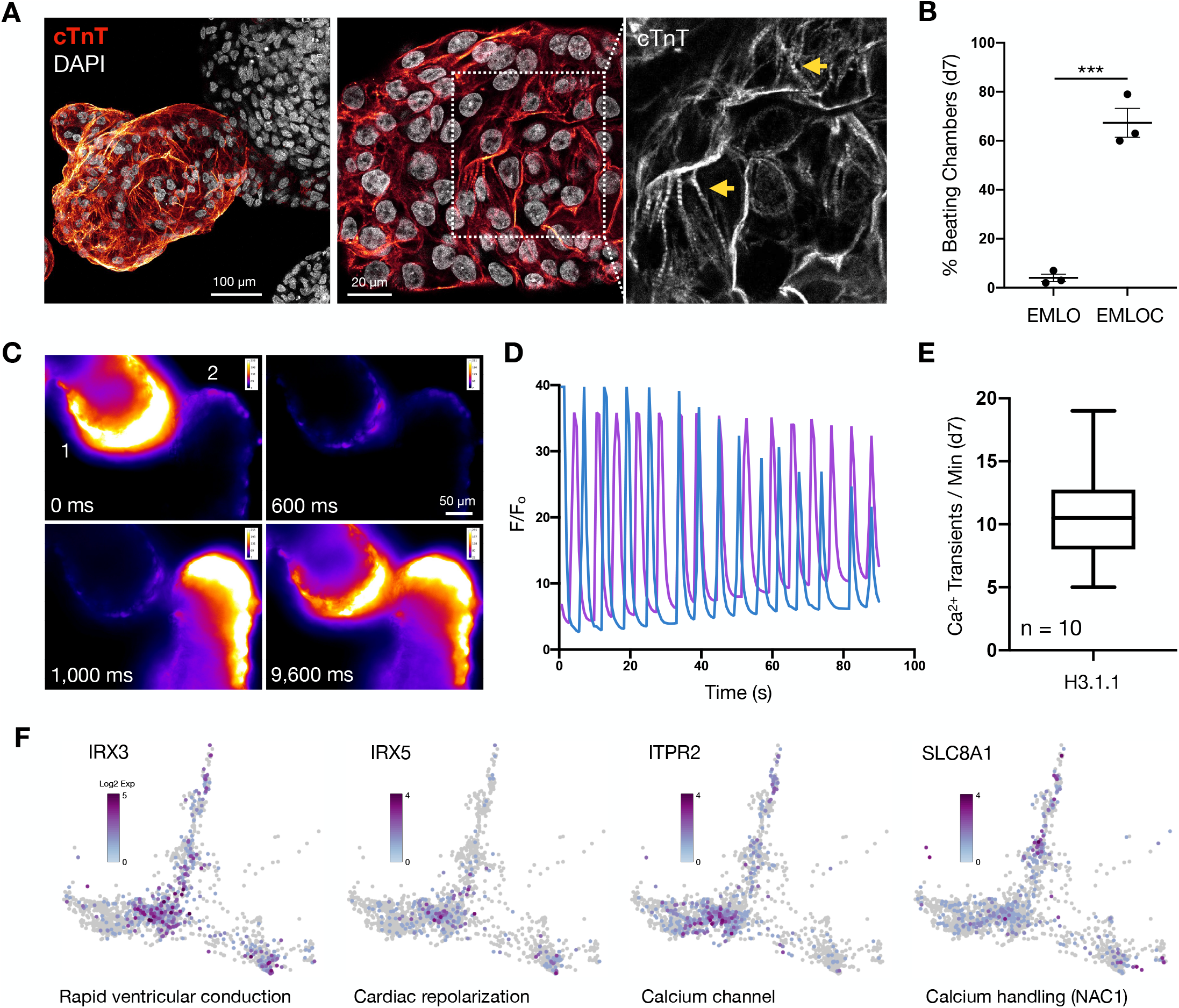
Spontaneous contractility and calcium handling in the EMLOC cardiac region. (**A**) Sarcomeres are visible with cTnT at day 7 in EMLOC formation. Progressive zoom with high magnification black and white image (right). Yellow arrows depict striations. (**B**) Percentage of EMLOCs with beating chambers versus trunk biased gastruloids formed by the original EMLO protocol (Olmsted and Paluh, 2021a, 2021b). (N = 3 repeat experiments; ***p = 0.0005, t = 10.40 df = 4 by unpaired two-tailed t-test). (**C**) Fluo-4 AM calcium imaging time course in two adjacent H3.1.1 EMLOCs. Individual scale bars provided. Images shown with fire LUT (ImageJ) and calibration bar. Individual scale bars provided. (**D**) Quantified F/F_o_ time series from (**C**). The two EMLOCs shown were captured in the same field and are representative of the population. (**E**) Box-and-whisker plot quantification of Fluo-4 AM Ca^2+^ transients/min in H3.1.1 day 7 EMLOCs (n = 10 EMLOCs, max = 19, min = 5, median = 10.5, q1 = 8, q3 = 12.75) from N = 2 separate differentiations. (**F**) Genes involved in rapid ventricular conduction (*IRX3*), repolarization (*IRX5*), calcium flux (*ITPR2*) and handling (*SLC8A1*/NAC1) are upregulated in EMLOCs. Day 16 scRNAseq data visualized by PHATE.

### Recapitulating early morphogenesis events in human developmental cardiac EMLOC gastruloids

The first cardiogenic structure to form in the anterior aspect of the mammalian embryo is called the cardiac crescent, which fuses to form the transversal heart tube that seals dorsally to generate a closed tube with outflow tracts (**Figure 4A**). We identified cardiac crescent-like structures at high penetrance in EMLOC (73.0 +/- 7.2%, mean +/- s.e.m.) versus EMLO generated gastruloids (2.7 +/- 1.8%; N = 3 repeat experiments; ***p = 0.0007, t = 9.474, df = 4 by unpaired two-tailed t-test) (**Figure 4B**). Cardiac crescent regions at day 4 in EMLOC formation contained cTnT+ cardiomyocyte progenitors co-localized with GATA6, a transcription factor required for high fidelity cardiogenesis (**Figures 4C and 4E**). The cardiogenic region increased in size with time, extending laterally away from the main body of the gastruloid (**Figure 4D**). The same region developed a cell-free interior, resembling early heart tube formation (**Figures 4A and 4F**) along with cardiac chamber morphological precursors (**Figure 4G**).

**Figure 4.**
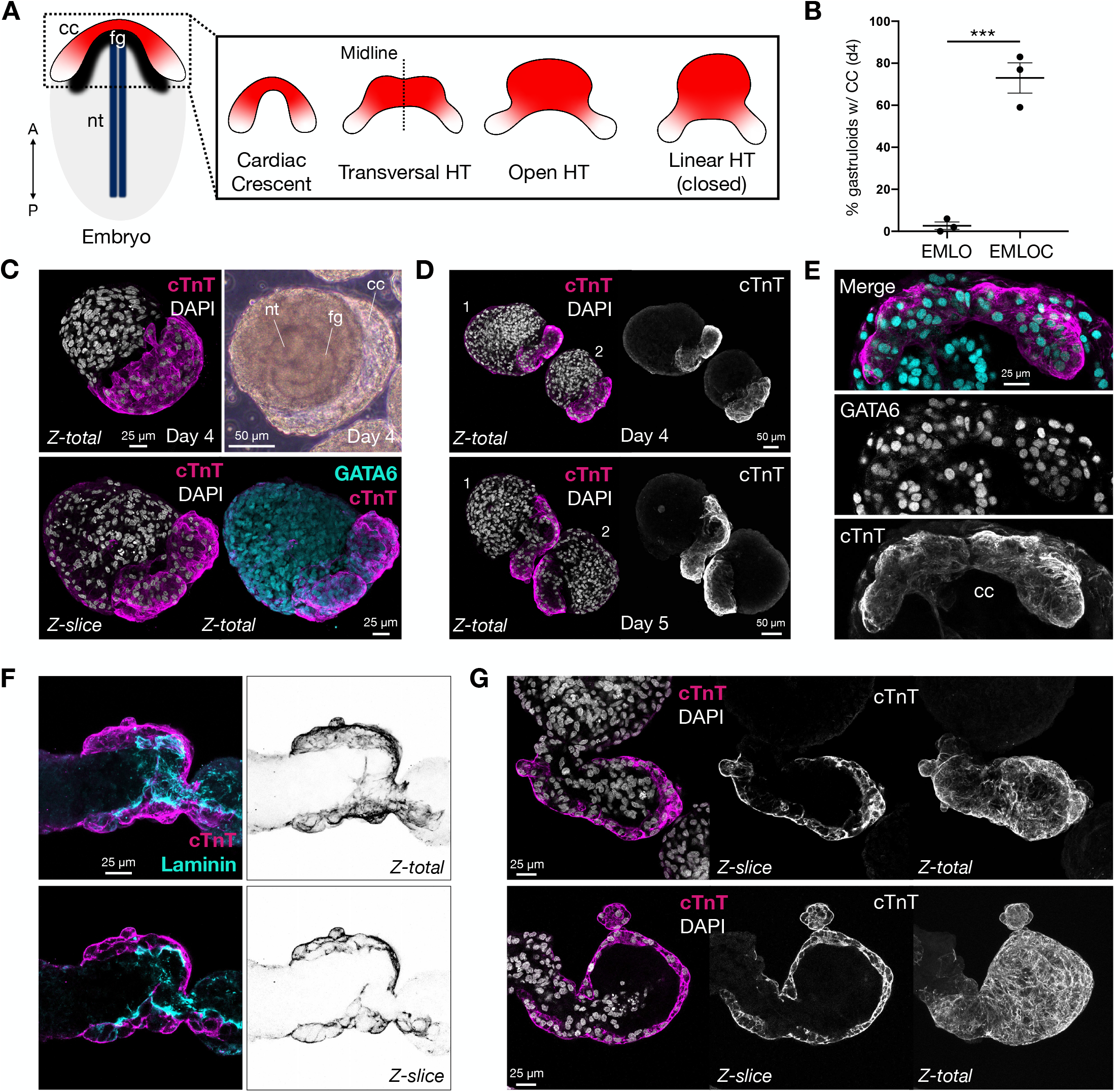
EMLOCs recapitulate early polarized heart tube formation events. (**A**) Cartoon schematic of the embryo depicting anterior cardiac crescent (cc, red/white), foregut cavity (fg, black), and neural tube (nt, dark blue). Anterior-posterior axis is indicated. The bilateral cardiac crescent fuses to form the transversal heart tube (HT), dorsally open heart tube, and linear heart tube (closed). (**B**) Percentage of day 4 gastruloids with cardiac crescent using the original EMLO protocol (Olmsted and Paluh, 2021a, 2021b) versus the optimized EMLOC protocol (N = 3 repeat experiments; ***p = 0.0007, t = 9.474, df = 4 by unpaired two-tailed t-test). (**C**) Day 4 EMLOCs exhibit cTnT+ cardiac crescent-like structures (magenta, top) with GATA6+ nuclei (bottom, cyan). Phase image of day 4 EMLOC is shown with labeled nt, fg, and cc corresponding structures (top right). (**D**) Comparison of cTnT+ cardiogenic regions in day 4 versus day 5 EMLOCs. Two adjacent EMLOCs are shown for each time point. (**E**) High magnification of cardiac crescent structure with cTnT (magenta) and GATA6 (cyan). Individual channels are shown without pseudocolor. (**F**) Immunofluorescence of cTnT (magenta) and laminin (cyan) with inverted cTnT channels depicts developing heart tube-like structure in EMLOCs (day 6). Z-slices and cTnT maximal projections are shown. (**G**) Cardiac chamber precursors in two separate EMLOCs (day 7). Z-slices and cTnT maximal projections are shown. Individual scale bars provided.

### EMLOC gastruloid cardiac morphogenesis occurs anterior to primitive gut tube endoderm

The developing anterior foregut derived from endoderm has been shown to be essential for cardiogenesis in multiple organisms through crosstalk with splanchnic mesoderm and by providing mechanical cues (**Figure 5A**) (Nascone and Mercola, 1995; Varner and Taber, 2012; Anderson et al., 2016; Kidokoro et al., 2018; Han et al., 2020). We previously demonstrated that the primitive gut tube can self-organize reproducibly in the EMLO gastruloid system (Olmsted and Paluh, 2020a). Therefore, we investigated whether this structure is present in the EMLOC gastruloids optimized for cardiogenesis. In mouse, this occurs anterior to the gut tube, including cuboidal epithelialization that is required for second heart field contributions to heart tube formation (Ivanovitch et al., 2017; Cortes et al., 2018). The splanchnic mesoderm is an established and reproducible feature of EMLO and EMLOC formation and also contributes to the gut tube (**Figure S3A and S3B**) (Olmsted and Paluh, 2021a). Using scRNAseq, a population of cells with anterior foregut identity (*FOXA2, NKX2-1, SHH, EPCAM*) was identified and clustered adjacent to the developing cardiac region (**Figure 2A**). Spatially organized FOXA2+ cells adjacent to the cardiac region were also identified using immunofluorescence. By normalizing fluorescence and gastruloid end-to-end length, an average profile for cTnT (cardiac) and FOXA2 (gut tube endoderm) was generated over the anterior-posterior axis to depict relative positioning in day 4 EMLOCs (**Figure 5B**) (N = 5 EMLOCs). Representative cTnT/FOXA2 immunofluorescence Z-slices are provided for day 5 and day 6 EMLOCs, in addition to CDH1 (E-Cadherin) and GATA6/Type 1 Collagen (**Figures 5C and 5D**). These data prioritize visualization of gut tube positioning relative to the cardiogenic region, demonstrating the appropriate posterior embryological trunk spatial organization with respect to cardiogenesis. We performed immunofluorescence imaging with cTnT and the proliferation marker Ki67, as well as CDH2 that is essential for ventricular wall morphogenesis (Miao et al., 2019). The co-immunofluorescence of the CDH2 biomarker with cTnT further revealed early organization of cells in the cardiac crescent into epithelial-like cytoarchitectures that is a contributing factor in heart tube formation (**Figure 5E**) (Cortes et al., 2018). This pattern was also observed by scRNAseq co-expression patterns of *CDH2* with ventricular biomarkers (**Figure S3B**).

**Figure 5.**
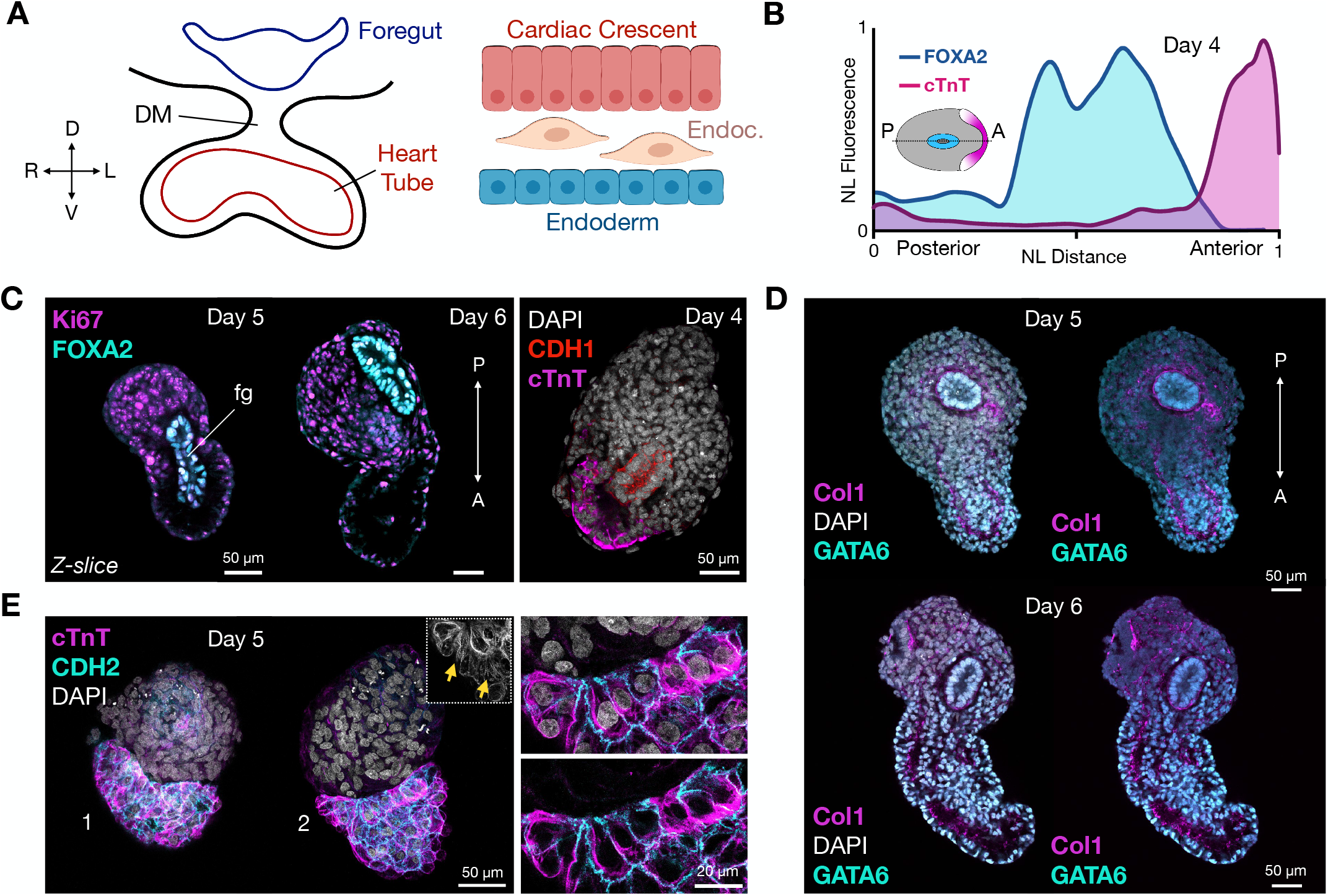
Cardiogenesis occurs anterior to gut tube endoderm. (**A**) Left: cartoon schematic of foregut and heart tube developmental cross-section with dorsal mesocardium (DM). The dorsal-ventral (D-V) and right-left (R-L) embryonic axes are shown. Right: cardiac crescent differentiation microenvironment. Cardiomyocyte progenitors (red), endocardium (tan), and definitive endoderm (blue) are shown. (**B**) Biomarker distribution using cTnT (cardiac, pink) and FOXA2 (gut tube, blue) in day 4 EMLOC gastruloids provided as smoothed curves. Normalized fluorescence was plotted over the anterior-posterior (A-P) axis normalized distance. (**C**) Left: Immunofluorescence Z-slices of FOXA2 (cyan)/Ki67 (magenta) depicts primitive foregut (fg) relative to cardiogenic region in day 5 versus day 6 EMLOC gastruloids. Right: CDH1/E-Cadherin depicts primitive gut tube (red) positioning relative to cTnT+ cardiogenic region (magenta). Cells are counterstained with DAPI. (**D**) Immunofluorescence Z-slices of GATA6 (cyan)/type 1 collagen (Col1, magenta) depicts primitive gut tube relative to cardiogenic region during chamber precursor formation in day 5 (left) versus day 6 (right) EMLOC gastruloids. (**E**) cTnT (magenta) and CDH2/N-Cadherin (cyan) co-localization in cardiac crescent reveals epithelization of cardiomyocyte progenitors. Inset depicts cTnT+ without pseudocolor. Yellow arrows depict striations. High magnification images with and without DAPI are provided (right) at the boundary of the EMLOC cardiogenic region. Individual scale bars provided.

We further investigated whether EMLOCs recapitulate distinct phases of cardiomyocyte differentiation and morphogenesis that are ongoing developmentally (**Figures S5**). As done *in vivo* (Ivanovitch et al., 2017), cardiomyocyte shape was used to indirectly infer distinct phases of cardiomyocyte differentiation versus structural morphogenesis with cellular proliferation. We characterized cardiomyocyte cell shape as rounded (morphogenesis and proliferation; **Figure S5A** top) versus adhesive flat/mosaic (differentiation; **Figure S5A** bottom) and quantified the relative proportion of EMLOCs exhibiting one phenotype or the other in single day 7 fixed samples (**Figure S5B**) (N = 4 replicates; 34 +/- 18% round, range 16 to 59%; 66 +/- 18% flat/mosaic, mean +/- s.e.m., range 41 to 84%; n.s. p = 0.1776, t = 1.754, df = 3 by paired two-tailed t-test). Together, these data demonstrate that features of *in vivo* cardiogenesis can be modeled in EMLOCs within the appropriate multi-lineage gastruloid microenvironment.

### EMLOCs exhibit specialization over heart tube length and multi-layering of chamber walls during morphogenesis

As the cardiac crescent is remodeled into the contractile primitive heart tube *in vivo*, specialization over the length of the tube establishes the future blueprints for the adult heart in terms of septated chambers and outflow tracts that transmit and receive blood (**Figure 6A**). Divisions of the embryonic heart are separated by minor constrictions in the tube. We identified day 7 EMLOCs with cardiogenic compartments resembling this stage in heart tube development (**Figure 6B**). After day 7, constricted tubes became dilated and had early divisions between chamber precursors (**Figures 6C and 6D**), visualized by 3D reconstructions and multi-dimensional analysis. The fluid-filled contractile cavities were completely surrounded by continuous cTnT+ cardiomyocytes indicative of myocardium. In addition, the cavities had open channels communicating with the posterior EMLOC compartment (**Figures 6C and 7D, Figure 7**). Genes involved in left-right asymmetry specification during *in vivo* cardiogenesis were also upregulated in the day 16 scRNAseq data set (*IRX3, HAND1, PITX2, RTTN*).

**Figure 6.**
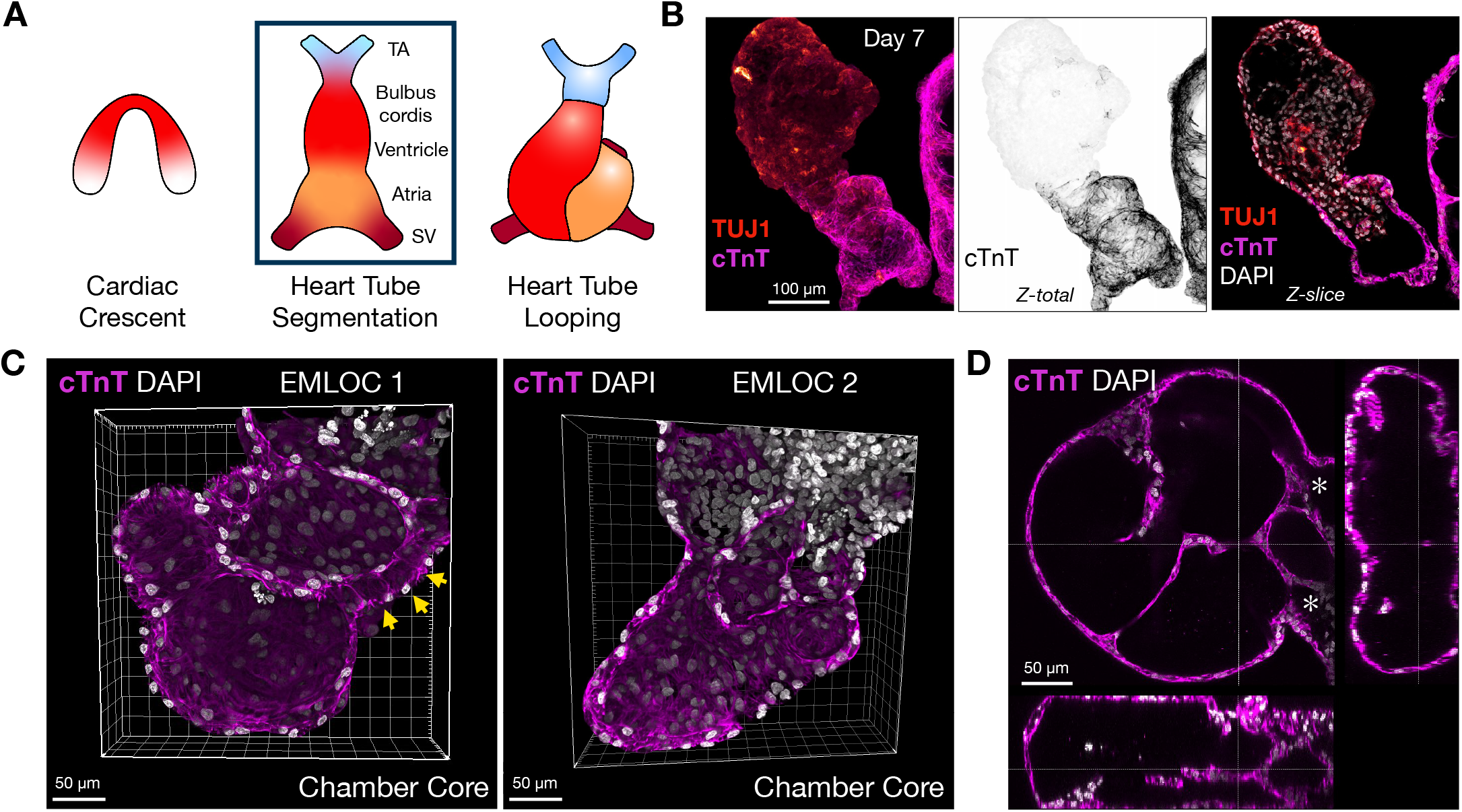
Myocardial expansion and heart tube morphogenetic specialization. (**A**) Cartoon schematic of cardiogenesis from cardiac crescent to heart tube specialization and heart tube looping. Truncus arteriosus (TA), sinus venosus (SV). (**B**) Heart tube staging in day 7 EMLOC gastruloids by cTnT (magenta) and TUJ1 (red), corresponding to the boxed stage in (**A**). Maximally projected Z-stacks and single Z-slice is shown. (**C**) 3D reconstruction of the anterior cardiac region with cTnT (magenta) and DAPI (grey) depicting putative outflow tract (yellow arrows) and chambers in two EMLOC gastruloids. (**D**) Multi-dimensional visualization of cTnT (magenta) and DAPI (grey). Sagittal and transverse planes are shown. Individual scale bars provided. Asterisks (*) indicate communication with proximal EMLOC compartment. Individual scale bars provided.

**Figure 7.**
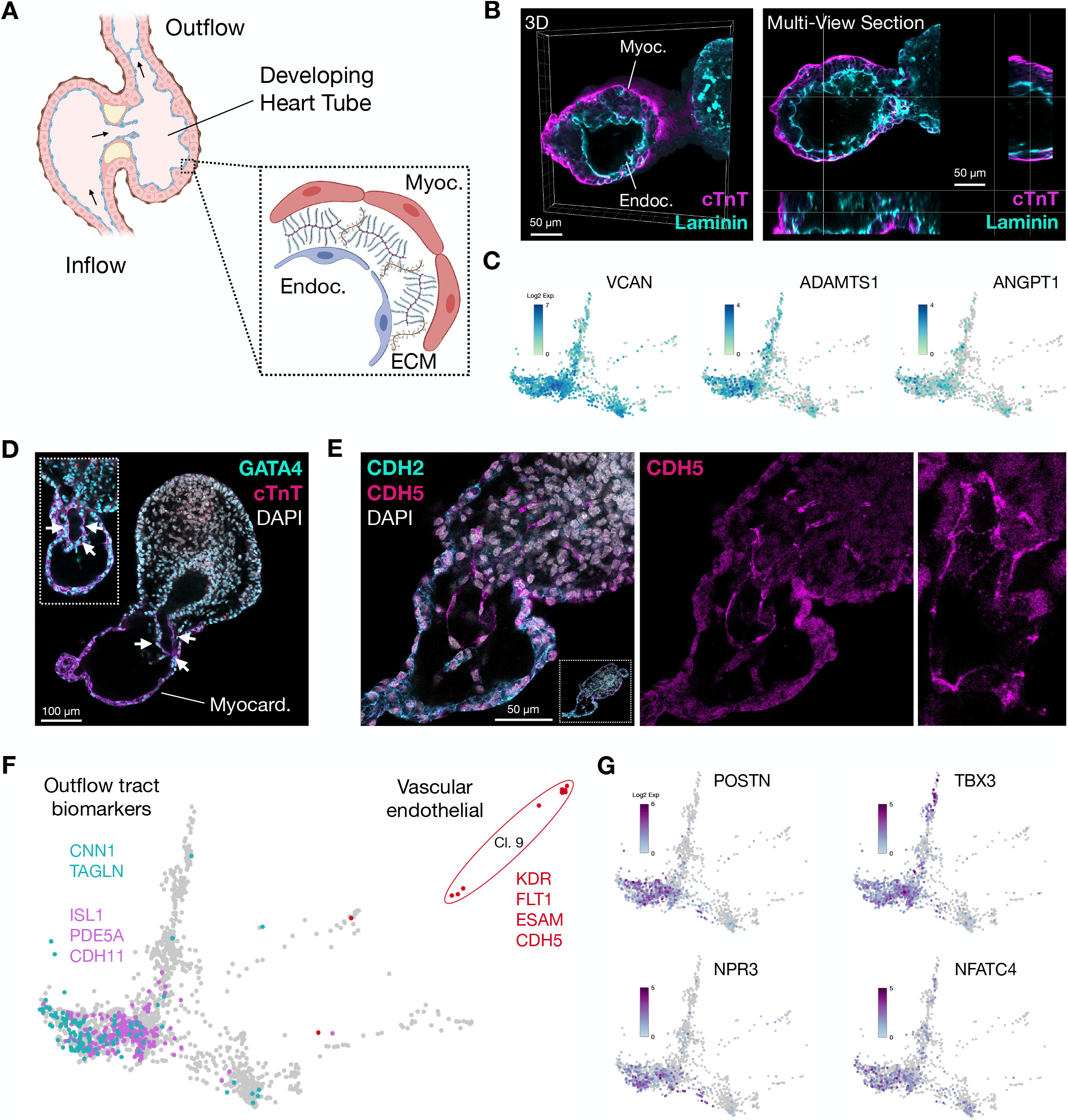
Chamber precursor morphogenesis in EMLOCs. (**A**) Cartoon schematic of a developing heart tube. Chamber wall layers are expanded to depict myocardium, extracellular matrix-rich cardiac jelly, and endocardium interior lining. (**B**) 3D reconstruction of the anterior cardiac chamber-like structures with cTnT+ myocardium (magenta) and Laminin+ interior (top-left). Single Z-slice multi-dimensional view of chamber (top-right). High magnification images are provided below. (**C**) *VCAN, ADAMTS1, ANGPT1* genes involved in cardiac jelly and its spatiotemporal degradation in day 16 EMLOC by scRNAseq, visualized using PHATE. (**D**) Immunofluorescence of cTnT, GATA4 and DAPI demonstrating putative outflow tract (white arrows) in two day 7 EMLOCs. Z-slice inset provides a second example. (**E**) Immunofluorescence of N-Cadherin (CDH2, cyan), VE-Cadherin (vascular endothelial cadherin/CD144/CDH5, magenta), and DAPI (grey) depicting endothelial biomarker expression lining the putative outflow tract. Rightmost zoom image depicts valve-like crescent structures. Individual scale bars provided. (**F**) Biomarkers for smooth muscle cells (*CNN1/TAGLN*) and the outflow tract (*ISL1/PDE5A/CDH11*) along with vascular endothelial cells (*KDR/FLT1/ESAM/CDH5*) (cluster 9). (**G**) *POSTN, TBX3, NPR3, NFATC4* genes involved in atrioventricular valve formation.

In developing heart chambers *in situ*, the chamber walls are multi-layered, with myocardium composed of working contractile and conducting cardiomyocytes comprising the outermost layer, and endocardium lining comprising the innermost layer (**Figure 7A**). An epicardial membrane surrounds these layers. Myocardium and endocardium are initially separated by ECM-rich “cardiac jelly” (Kim et al., 2018) that is degraded with time, necessary for chamber morphogenesis. Using 3D image reconstruction and multi-dimensional analysis, we show that chamber wall organization was recapitulated in developing EMLOC chamber-like structures (**Figure 7B**). We identified gene biomarkers of cardiac jelly ECM and its spatiotemporal degradation *VCAN*, *ADAMTS1*, and *ANGPT1* that were highly expressed in day 16 EMLOCs (**Figure 7C**) (Kim et al., 2018). Notably, an interior channel partially lined by cTnT+ cells was also identified with high penetrance, postulated here to be initiation of the putative outflow tract due to its appropriate positioning (**Figure 7D**) and that is lined by segmental CDH5 immunostaining extending out into the posterior compartment of the EMLOC (**Figure 7E**). By scRNAseq, we identified cells with combinations of biomarkers for smooth muscle (*CNN1/TAGLN*), outflow tract development (*ISL1/PDE5A/CDH11*), and well-differentiated vascular endothelium (cluster 9; *KDR/FLT1/ESAM/CDH5*) (**Figure 7F**). Nodal and valvar biomarkers were also present with a similar distribution (*POSTN/TBX3/NPR3/NFATC4*) (**Figure 7G**). These data, taken together, are consistent with cardiogenesis in EMLOCs proceeding in an appropriate spatiotemporal manner and detailed morphological and gene expression changes, according to aspects of in situ development.

### EMLOCs capture neurogenesis within a neuro-cardiac model of human trunk development

Our previous EMLO approach (Olmsted and Paluh, 2021a, 2021b) was developed to study early neurogenesis events in trunk development. To investigate early neural lineage biomarkers in EMLOCs we performed immunofluorescence and scRNAseq (**Figure 8**). TUJ1 immunostain was first identified in cells opposite the anterior cardiac domain with low level staining, typical of neural stem/progenitor cells that express this protein at lower levels (**Figure 8A**). The initial emergence of neurons at day 7, identified by morphology and biomarkers, parallels that seen for the original EMLO protocol (Olmsted and Paluh, 2021a, 2021b). In EMLOCs, the posterior region of neurogenesis emerged from one to several SOX2+/TUJ1+ neuroectodermal rosettes and the number of neurons increased significantly over time (**Figures 8B-8D**). Given the relatively low number of neurons present at the day 7 time point when spontaneous contractility is already occurring, it is unlikely that neuronal function plays a significant signaling role in initiating spontaneous cardiogenic contractions at this early stage. The increase in the number of TUJ1+ neurons in EMLOCs was quantified between days 7 and 18 (**Figure 8E**) (day 7: 5 +/- 2 TUJ1+ neurons, mean +/- s.e.m; day 18: 218 +/- 24 TUJ1+ neurons; ****p < 0.0001, t = 8.929, df = 18 by unpaired two-tailed t-test). We also quantified the proportion of EMLOCs with a neuronally integrated cardiac compartment between days 7 and 25 (**Figure 8F**), which similarly increased with time (day 7: 3.3 +/- 1.9% of population; day 25: 55.8 +/- 2.9; **p = 0.0011, t = 12.58, df = 3 by paired two-tailed t-test; N = 4 replicate experiments). A nidus of neurogenesis from neural rosettes occurred within GATA6+ surrounding mesenchymal-like tissue (**Figure 8G**). Notably, the gut tube can be distinguished from surrounding rosettes by a laminated acellular border, whereas the rosettes are more continuous with the adjacent GATA6+ cells (**Figure S6**). In EMLOCs, the open channels from the cardiac chambered region connect with the proximal compartment containing neural rosettes.

**Figure 8.**
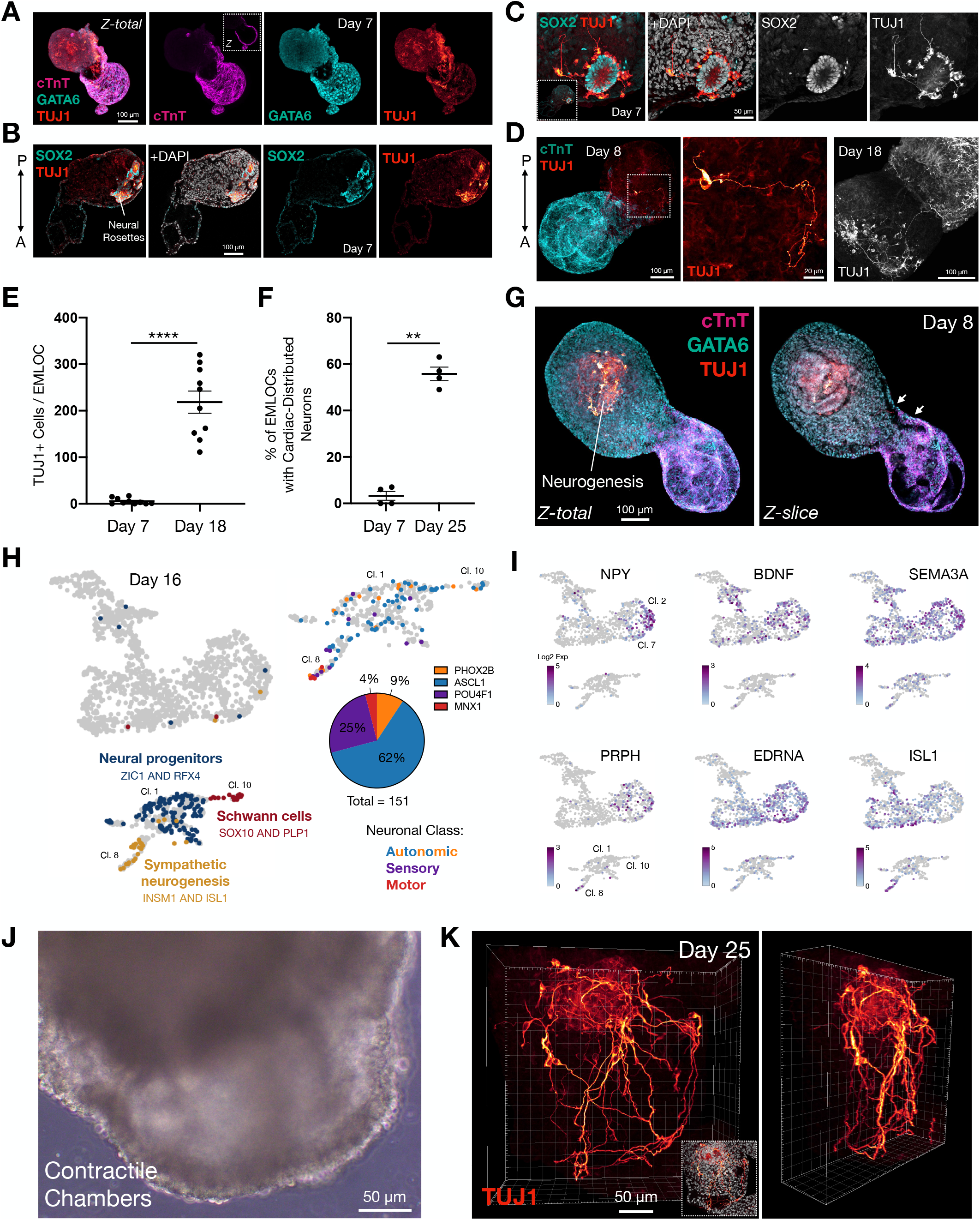
Neuron co-development and population of the cardiac region. (**A**) Neural rosette biomarkers SOX2 (cyan) and TUJ1 (red) emerging posteriorly in day 7 H3.1.1 EMLOC, counterstained with DAPI. (**B, C**) SOX2+ rosette and neurogenesis (TUJ1) counterstained with DAPI along with high magnification (**C**). Inset is whole EMLOC. (**D**) Immunofluorescence of cTnT (cyan) and TUJ1 (red) depicting single neuron in day 8 EMLOC (left, white dotted box) and zoom in (middle panel). Comparison with TUJ1+ neuron tract in day 18 EMLOC (far right panel). (**E**) Absolute number of TUJ1+ cells in day 7 versus day 18 EMLOCs (n = 10 per time point). ****p < 0.0001, t = 8.929, df = 18 by unpaired two-tailed t-test. (**F**) Proportion of EMLOCs with TUJ1+ neuronal fibers distributed within the cardiac region at day 7 versus day 18 (N = 4 replicate experiments; day 7: 3.3 +/- 1.9% of population; day 25: 55.8 +/- 2.9; **p = 0.0011, t = 12.58, df = 3 by paired two-tailed t-test). (**G**) Nidus of neurogenesis posterior to gut tube endodermal cells and cardiogenic region in day 8 EMLOC by TUJ1 (red), GATA6 (cyan), and cTnT (magenta). Maximally projected Z-stack (Z-total) and single Z-slice shown. White arrows point to communicating channels. Gut tube endoderm is laminated while neural rosettes are continuous with surrounding GATA6+ cells. (**H**) Left: UMAP representation of day 16 EMLOC scRNAseq highlighting trunk neural progenitors (*ZIC1/RFX4*), sympathetic neurogenesis (*INSM1/ISL1*) and Schwann cell glia (*SOX10/PLP1*). Right: quantification of neuronal class proportions in clusters 1, 8, 10 as autonomic (*ASCL1* 62%*, PHOX2B* 9%), sensory (*POU4F1* 25%) and motor (*MNX1* 4%). (**I**) Genes in the cardiogenic region involved in neuronal patterning and innervation. *NPY, BDNF, SEMA3A, PRPH, EDNRA, ISL1* shown by UMAP (day 16). (**J**) Phase contrast image of contractile EMLOC chamber-like structures (see also **Movie S5**). (**K**) 3D reconstruction of TUJ1+ neuronal fiber intercalation with chamber-like myocardium in day 25 EMLOC (see also **Movie S6**). Rotated view is shown. Individual scale bars provided.

In the integrated scRNAseq dataset, clusters 1, 8 and 10 predominantly represented the neural lineage. Neural progenitors (cluster 1; *ZIC1, RFX4, HES5, FABP7, EDNRB, NTRK2, OLIG3, MSX1*) became specialized neuronal subtypes (cluster 8; *INSM1, ELAVL3, DLG4, CAMK2A, SLC18A3, SLC17A6, CHRNA3, NTRK3*), and a population of neural crest-derived Schwann cells was present (cluster 10; *SOX10, PLP1, MPZ, S100B, TFAP2B, NGFR*) (**Figures 1D-1F**, **Figure 8H**). Combined *INSM1/ISL1* expression indicates that sympathetic neurogenesis occurs in EMLOCs, which is particularly relevant to developing cardiac innervation. Restricted expression of *HOXC6* and *HOXC9* to these clusters supports spinal cord and trunk identity. Schwann cells increased ∼16-fold from the day 7 to day 16 time point. As well, biomarkers of specialized neuronal subtypes that were lacking at day 7 began to emerge. The transcriptional phenotype for differentiating autonomic neurons that is *ASCL1* (93/151 neurons, ∼62%) and *PHOX2B* (14/151 neurons, ∼9%) predominated versus sensory neurons *POU4F1*/BRN3A (38/151 neurons, ∼25%) or motor neurons *MNX1*/HB9 (6/151 neurons, ∼4%) (**Figure 8H**). This finding is distinct from EMLOs (Olmsted and Paluh, 2021a), in which motor neurons were primarily generated.

### EMLOCs express biomarkers of cardiac innervation

In parallel with neurogenesis, axonal projections navigating the extracellular space to their target sites are expected to require spatial signals to generate the selective patterning on organs for innervation. Molecular and morphogenic features of the developing heart must therefore play an active role in establishing autonomic innervation, where the proper cellular milieu and receptive fields for innervation will dictate selective neuronal interactions. As such, we identified several genes with known roles in this process that were expressed in the cardiogenic region of the UMAP plot including and that code for neuropeptide Y (*NPY*), brain-derived neurotrophic factor (*BDNF*), semaphorin 3A (*SEMA3A*), peripherin (*PRPH*), endothelin receptor type A (*EDNRA*), and ISL-1 (**Figure 8I**). Genes involved in autonomic neurogenesis and cardiogenesis such as *ISL1* also play a role in development and innervation of cardiac pacemaker cells that dictate automaticity and participate in the conduction system apparatus.

By immunofluorescence, neurons were not identified within the cardiac region at the earlier day 7 and day 8 time points. We therefore analyzed the degree to which cardiogenic and neurogenic regions of the EMLOCs co-develop and integrate. In EMLOCs at day 16 or more in formation, neurons were observed both in the posterior compartment and intercalated with cardiomyocytes anteriorly, resembling *in vivo* ganglionated plexuses that characterize heart innervation (Ashton et al., 2018) (**Figure S7**). In day 25 EMLOCs (**Figures 8J and 8K**), the cardiogenic region remained contractile (**Figure 8J, Movie S5**) and neurons formed elaborate structural networks within the myocardium that are excluded from the chambers, appreciated using 3D reconstruction analysis (**Figure 8K, Movie S6**). Terminating neuronal fibers on cardiomyocytes were identified in part as axons using the phospho-tau (Ser214) immunostain (**Figure S7E**). We anticipate that the co-habitation of neurons and cardiomyocytes in the same region is a precursor to neuro-cardiac cooperative interactions such as initiation of innervation.

## DISCUSSION

The consistent lifelong critical functioning of the adult human heart is established during embryonic development in a process known as cardiogenesis. EMLOCs provide the first detailed insights into integrated neurogenesis and cardiogenesis in a human gastruloid developmental model. The complex process of cardiogenesis requires short-range interactions with surrounding tissues and occurs in conjunction with long-range input by neurons through progressive innervation (Harvey, 2002; Hasan, 2013). As the first organ to function in the embryo, the developing heart begins to supply blood to the growing fetal brain as a closed tube, even before undergoing dramatic structural reorganization and maturation into septated chambers with outflow tracts. Such complexity and dependence on multiple non-cardiac tissue inputs has made it difficult to recapitulate human heart development using traditional *in vitro* models, requiring instead refined gastruloid technologies.

The intracardiac nervous system is sometimes colloquially referred to as the “brain within the heart” (Campos et al., 2018). Using sophisticated methodologies such as optogenetic stimulation, the role of peripheral cardiac neural circuitry in pacemaking and conduction is beginning to be understood (Rajendran et al., 2019; Fedele and Brand, 2020). Innervation of the heart *in vivo* is predominately autonomic, where sympathetic neurons can directly innervate working cardiomyocytes in the ventricular wall, and are networked as so-called ganglionated plexuses (Zaglia et al., 2017). The neurons that begin to develop in EMLOCs at the time when spontaneous contractility is first observed (∼day 7) are unlikely to substantially contribute to contractile function, since at this stage they are relatively few in number and do not project into the cardiogenic region. This is consistent with *in vivo* development where contractility of the heart tube occurs prior to innervation that is established later (George et al., 2020). However, organized neuronal networks resembling ganglionated plexuses were observed as EMLOCs progressively matured. At the day 7 time point, neurons were localized distantly from the cardiogenic region, before expanding significantly in number to migrate, embrace and populate the myocardial layer over time. One potential explanation is a microenvironment switch from axon-repulsive to axon-permissive as ECM in the cardiac jelly is degraded. The ECM-rich cardiac jelly *in vivo* contains chondroitin sulfate proteoglycans and other components known to exert repulsive or pausing effects on axons during navigation and regeneration (Tom et al., 2004). Degradation of the cardiac jelly during development is physiologic and required for normal cardiac chamber morphogenesis (Kim et al., 2018). Differential regulation of *SEMA3A* expression may also play a role. Within this framework, our data support the adherence of EMLOC events to physiologic spatiotemporal developmental processes for establishing contractile chambers with supplied neurons (Hasan, 2013; George et al., 2020). Neuromuscular interactions between cardiac innervating neurons and cardiomyocytes at the “neuro-cardiac junction” remains poorly understood (Zaglia et al., 2017) including biomarkers. Synapses with cardiomyocytes are postulated to be mediated through an alternate structure other than the nAChR machinery in skeletal neuromuscular junctions (Sargent and Garrett, 1995). Traditional 2D hiPSC differentiation protocols that generate human neurons and cardiomyocytes separately and then co-culture these cells to obtain structural and functional detail are typically used to study innervation, as has been done for the skeletal muscle neuromuscular junction (Darabid et al., 2014; Steinbeck et al., 2016). EMLOCs are expected to provide a developmental and spatiotemporal perspective of the neuro-cardiac junction.

Efforts to study heart development and function using human cells also focus on separate cardiac mechanisms and include combined tissue engineering platforms and solutions (Ma et al., 2015; Macqueen et al., 2018). Such top-down human intervention of biofabricated tissues and organs has not yet achieved developmentally patterned neuronal innervation (Das et al., 2020), but may benefit from this EMLOC study. Gastruloid models that more closely mimic embryogenesis are an exciting alternative to achieve and study organogenesis (van den Brink et al., 2014; Beccari et al., 2018; Moris et al., 2020). A recent study with mESCs made significant advances and achieved early key features of cardiogenesis (Rossi et al., 2021). Our developmental model of human cardiogenesis in gastruloids further advances cardiac models by including neuronal co-development and association with the myocardium. The scRNAseq analysis that indicates that we have established multiple prerequisites for innervation. In our previous study of EMLO gastruloids that generated CNS and PNS integration with mixed lineage trunk identity (Olmsted and Paluh, 2020a), we achieved self-organized spinal neurons, neural crest, and a primitive gut tube surrounded by splanchnic mesenchyme, thereby providing much of the ideal cardiogenic microenvironment. By modifying the EMLO protocol (Olmsted and Paluh, 2020b) to include pro-cardiogenic and angiogenic factors, VEGF and ascorbic acid, that were applied in the mESC *in vitro* cardiogenesis study (Rossi et al., 2021), we achieved coupled cardiogenesis and neurogenesis. Through comprehensive biomarker analysis and live cell calcium imaging, we demonstrate here that EMLOCs recapitulate numerous key features of human cardiogenesis including cardiac crescent transformation into the contractile heart tube, cardiomyocyte differentiation versus remodeling phases, and formation of chamber- and outflow tract-like structures. Cardiogenesis occurs anterior to primitive gut tube-like endodermal cells that *in vivo* are thought to be required (Nascone and Mercola, 1995; Schultheiss et al., 1995; Varner and Taber, 2012; Anderson et al., 2016; Kidokoro et al., 2018; Han et al., 2020).

A limitation of this study is the use of one hiPSC line and ability to evaluate only two developmental time points by scRNAseq of the numerous stages analyzed and described. Nine lines previously evaluated for EMLO formation revealed reproducibility of structural organization and cell types but with differences in efficiency between lines that can be optimized (Olmsted and Paluh, 2021a; 2021b). A second focus is expected to address how refined developmental insights can be coupled with bioengineering technologies for development of 3D organogenesis platforms bringing in additional features of vascularization. To understand the establishment and function of the neuro-cardiac junction in EMLOCs, electrophysiologic and optogenetic characterization of multi-tissue function beyond calcium imaging of neurons, cardiomyocytes, and neuro-cardiac junction biomarkers is needed. Compared to the well-studied cholinergic neuromuscular junction between motor neurons and skeletal muscle, much less is known about the autonomic synaptic junctions between neurons and working cardiomyocytes including all relevant biomarkers necessary to investigate innervation. Spatiotemporal analysis of scRNAseq data is further benefiting human developmental cell atlantes to complement existing databases for adult tissues (The Human Protein Atlas), and is being pursued for human cardiogenesis (Asp et al., 2019; Cui et al., 2019).

We anticipate that EMLOCs will open new opportunities to study fundamental questions on neuromodulation of contracting cardiomyocytes with relevance to neurocardiogenic syncope and other neural-based arrhythmia pathologies (Ashton et al., 2018). As well, such a neuro-cardiac model system is expected to provide fundamental insights into the pathophysiology of congenital heart disease and potential treatments in addition to viral infection studies and *in vitro* pharmacotherapy testing and discovery. As a drastically needed component of *in vitro* stem cell systems, innervation in non-neural tissue, organ, and embryo models (Das et al., 2020; Sahu and Sharan, 2020) is beginning to be achieved in EMLO and EMLOC gastruloids to advance innervation research. We expect that this approach and model will have broad biomedical relevance for neuro-cardiac development and human organ innervation initiatives.

### Limitations of this Study

In this work we develop and optimize a gastruloid model enabling co-development and self-integration of human neuronal and cardiac tissue precursors in a multicellular, multi-lineage context. These results extend our previous work with neurogenesis and gut development in EMLO gastruloids to promote concomitant cardiogenesis that recapitulates multiple key features of *in vivo* heart development. In EMLOCs, neurons are produced endogenously in the context of the developing cardiac region as opposed to by separate differentiation and subsequent combination by fusion or in co-cultures. Genetic manipulation for optogenetic activation of neuro-cardiac pathways will inform on circuit formation and maturation. However, optical resolution in thick samples with rapidly paced signaling and cell-cell communication events requires sophisticated imaging techniques or further manipulation of the spatially organized cells such by slice cultures that can help to dissect assembled pathways. Additional single cell sequencing at multiple extended time points will be informative to clarify the diversity of cell types, transition states, and signaling events. Future studies are expected to focus on continued developmental progression, particularly addressing functional innervation, and the impact of these processes on diseases that include ethnic contributions. EMLOCs therefore enable an exciting frontier to more rapidly address neural innervation of non-neural structures in gastruloids that has remained a challenge and priority of the field, and will continue to benefit from emerging technologies and resources.

## ACKNOWLEDGEMENTS

This work was funded by SUNY Polytechnic SEED 917035-21 and CATN2 MIP awards. It used published lines developed and initially characterized through previous grants awarded to the Paluh laboratory for New York State stem cell research (NYSTEM) and spinal cord injury (NYSCIRB) research. Analysis by scRNAseq was performed at the SUNY Buffalo Genomics and Bioinformatics core (SBGB). Jonathan Bard (SBGB) assisted in assembling the integrated scRNAseq dataset and visualization methods by UMAP and PHATE for downstream analysis.

## AUTHOR CONTRIBUTIONS

J.P. and Z.O. conceived of the project and experimental design and analyzed data and co-wrote the manuscript. Z.O. performed EMLOC formation and characterization experiments, and composed figures.

## INCLUSION AND DIVERSITY

Dr. Paluh received previous grant support (NYSTEM) to generate hiPSC lines with increased ethnically diverse representation for basic and clinical science, including the human Hispanic Latino iPSC line H3.1.1 used in this study. The authors have previously published several studies that also apply hiPSC lines from self-reported African American, Hispanic Latino, and Asian American fibroblast donors.

## SUPPLEMENTAL MOVIES

**Movie S1. Contractile EMLOCs in suspension culture at day 7**. Phase contrast real-time movie at 5x magnification. Related to **Figure S1**.

**Movie S2. Contractile EMLOCs in suspension culture at day 7**. Phase contrast real-time movie at 20x magnification. Related to **Figure S1**.

**Movie S3. Live cell calcium imaging for contractility with whole EMLOCs at day 7**. Example 1 with Fluo-4 AM dye. The cardiogenic regions of the two EMLOCs are adjacent. 200 ms interval, 50 ms exposure time, 17 frames per second. Related to Figures 3C**-3E**.

**Movie S4. Live cell calcium imaging for contractility with whole EMLOCs at day 7**. Example 2 with Fluo-4 AM dye. Calcium fluorescence correlates with visible contractility. 200 ms interval, 50 ms exposure time, 17 frames per second. Related to Figures 3C**-3E**.

**Movie S5. Contractile EMLOCs in suspension culture at day 25**. Phase contrast real-time movie at 20x magnification. Related to Figure 8J.

**Movie S6. 3D reconstruction of innervated cardiac chambers**. Generated using Imaris. Related to Figure 8K.

## METHODS

### RESOURCE AVAILABILITY

#### Lead contact

Further information and reasonable requests for resources should be directed to and will be fulfilled by the Lead Contact, Janet L. Paluh.

#### Materials availability

This study did not generate unique reagents.

#### Data and code availability

The scRNAseq data is available through GEO (GSE194356). The authors declare that all other data supporting the findings of this study are available within the article and its Supplementary Information files or from the corresponding author upon reasonable request.

### KEY RESOURCES TABLE

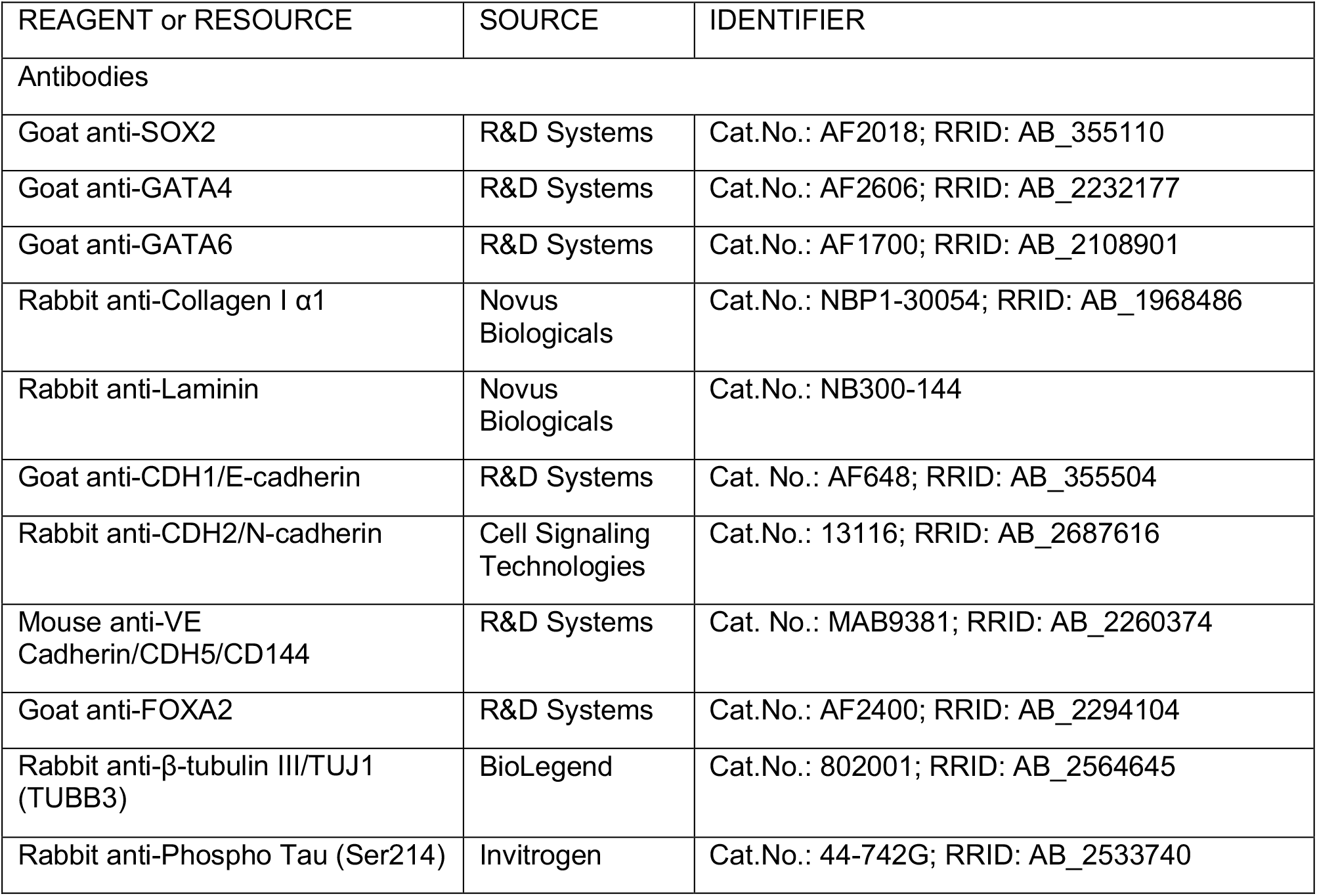
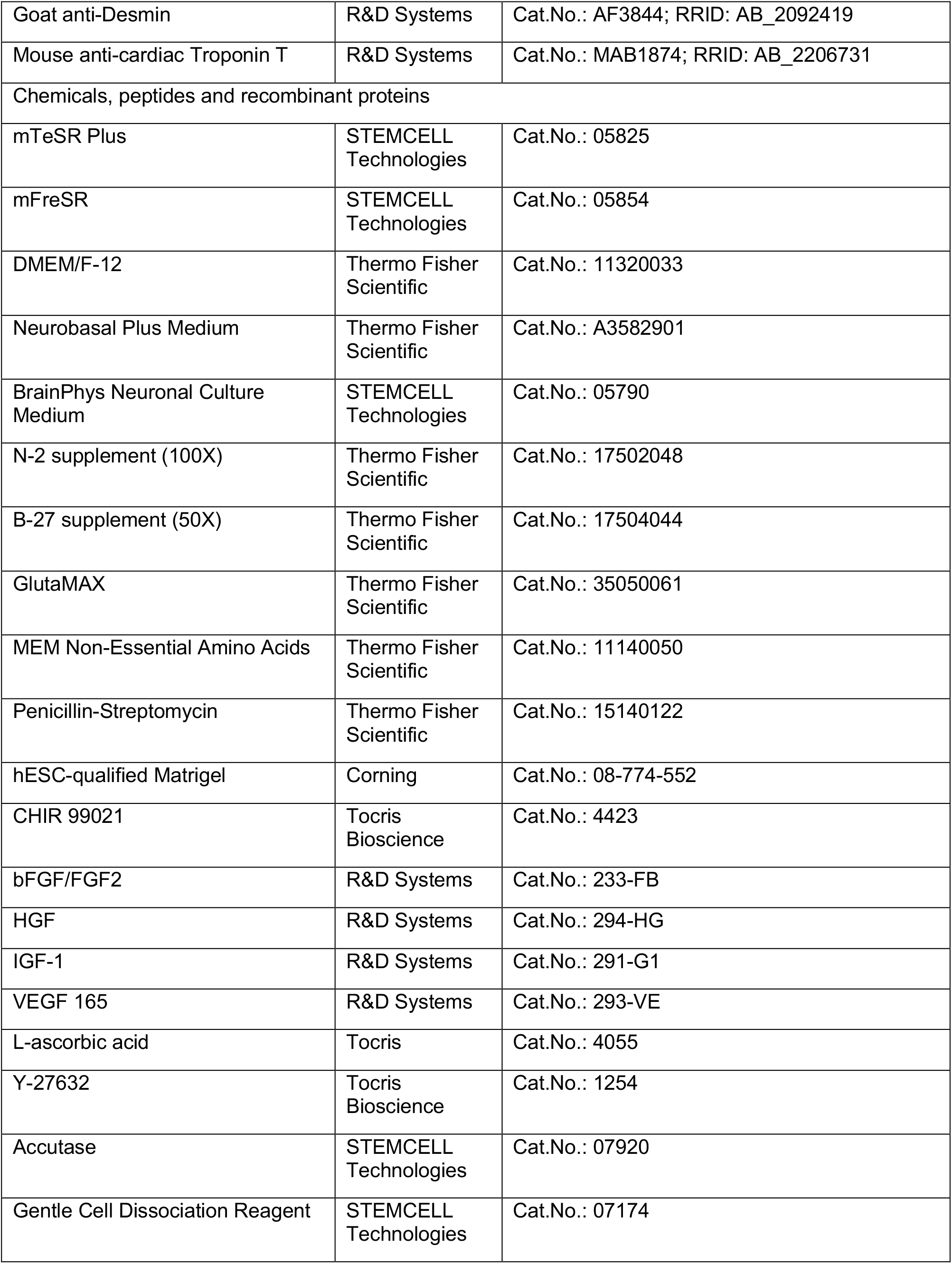
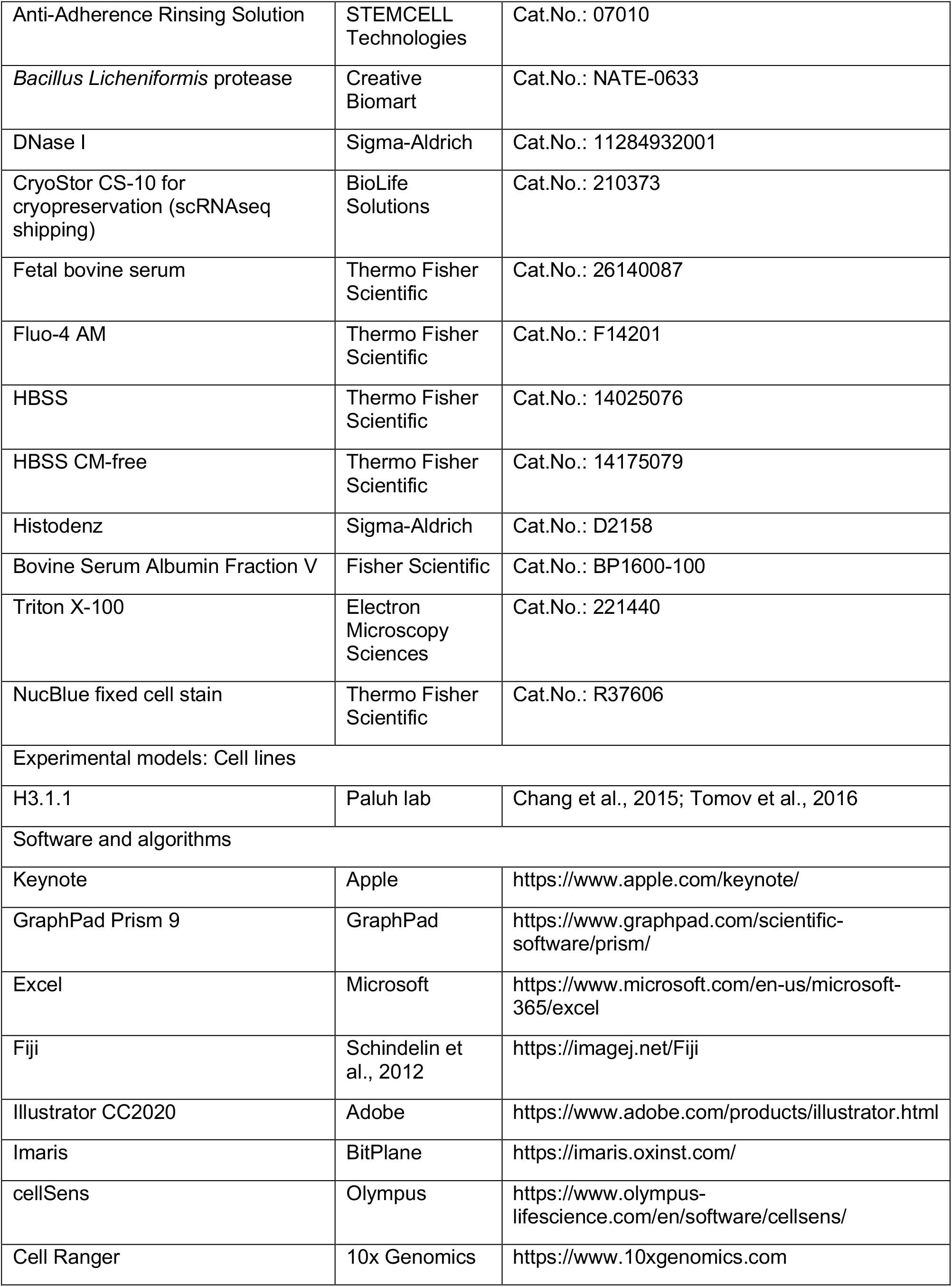
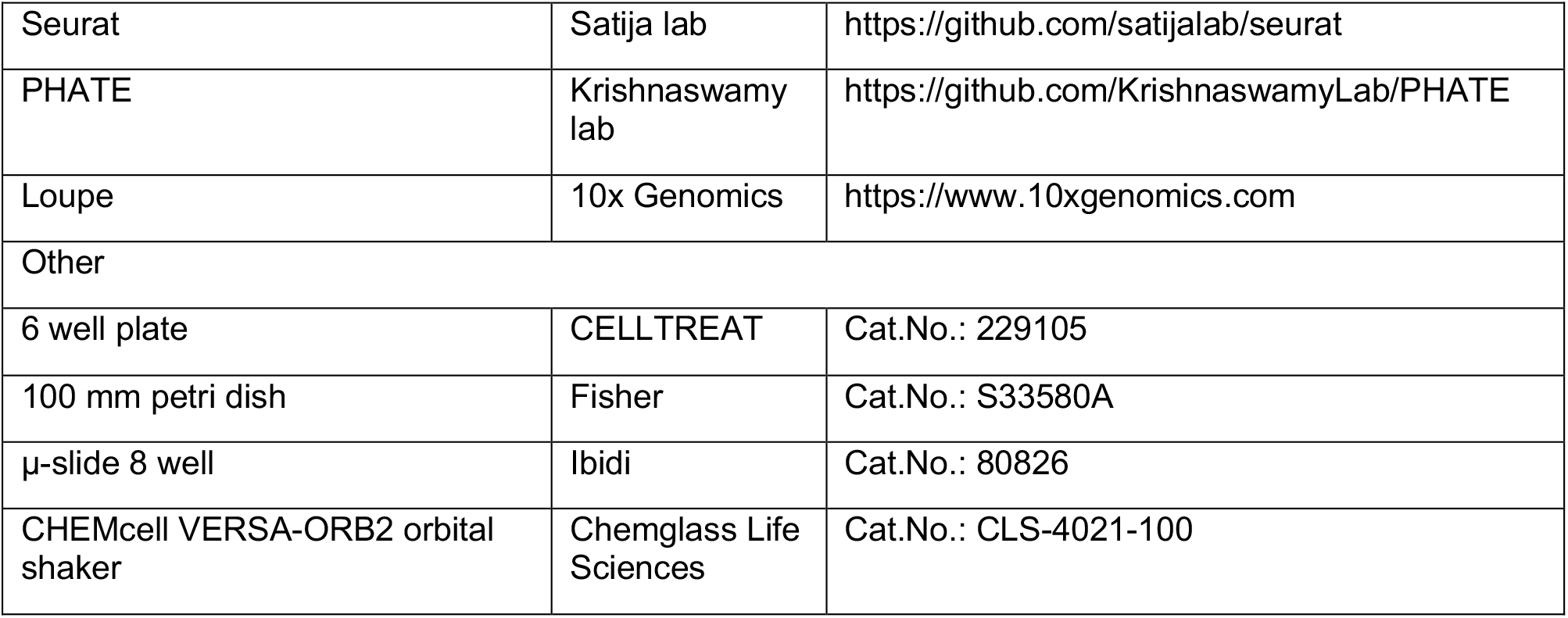

### EXPERIMENTAL MODELS AND SUBJECT DETAILS

#### Human induced pluripotent stem cells

We previously derived hiPSC lines from Coriell de-identified human fibroblast samples from consenting donors, including the Hispanic-Latino H3.1.1 line used in this study (Chang et al., 2015; Tomov et al 2016). Line H3.1.1 was reprogrammed with Yamanaka factors by the laboratories of Dr. Paluh and Dr. Jose Cibelli from these donor fibroblasts and comprehensively characterized for pluripotency (immunofluorescence, RT-PCR), G-band karyotype, teratoma formation, multi-lineage differentiation, bulk RNA-Seq, ChIP-Seq, and used in multiple studies from this lab. Recent G-band karyotype validation and pathogen analysis was performed by Cell Line Genetics, Inc. (NY, NY). H3.1.1 hiPSC colonies were cultured in mTeSR Plus supplemented with 1x penicillin-streptomycin (P-S) on hESC-qualified Matrigel (1:100 dilution; Corning) in a humidified incubator at 37°C, 5% CO_2_. Cultures were passaged 1:6 in 6-well plates every 4-7 days using Gentle Cell Dissociation Reagent (GCDR, STEMCELL Technologies). Cells were cryopreserved in mFreSR.

### METHOD DETAILS

#### EMLOC formation

EMLOs were formed similarly to as previously described (Olmsted and Paluh, 2021a, 2021b) with several important differences detailed as follows. H3.1.1 adherent hiPSC colonies were maintained in mTeSR Plus pluripotency medium as described above. At ∼60% confluency, pluripotency medium was changed to induction medium (N2B27 basal medium supplemented with 3 μM CHIR 99021, 40 ng/ml basic fibroblast growth factor FGF2). N2B27 basal medium: 1:1 DMEM/F-12:Neurobasal Plus medium, 2% (v/v) B27 Plus supplement, 1% (v/v) N2 supplement, 1x GlutaMAX, 1x MEM Non-Essential Amino Acids, 1x P-S. Adherent hiPSC colonies were induced for two days the one exchange of fresh medium at 24 h. On the day of aggregation, cells were dissociated with 1:1 Accutase:HBSS (Ca-Mg free) at 37°C for 5 min followed by manual trituration with a P-1000 pipette. Six-well plates were pre-treated with Anti-Adherence Rinsing Solution (STEMCELL Technologies) for 5 min incubation at room temperature followed by two rinses with equal volumes of HBSS. Cells were resuspended in N2B27 supplemented with 10 ng/ml FGF2, 2 ng/ml IGF-1, 2 ng/ml HGF (R&D Systems) and 50 μM Y-27632 (Tocris Bioscience). For aggregation, the single cell suspensions were added at a density of 2 x 10^6^ cells/ml (2 ml per well, 4 x 10^6^ total cells). Gastruloids were aggregated overnight using an orbital shaker at 75 rpm clockwise in a humidified incubator with 5% CO_2_. The next day, one-half volume of medium was replaced with fresh medium N2B27 supplemented with 4 ng/ml IGF- 1, 4 ng/ml HGF, 20 ng/ml FGF2 to maintain the same concentration of growth factors in the culture medium after one-half volume addition. At 48 h, the entire volume of medium was replaced with N2B27 basal medium supplemented with 5 ng/ml VEGF, 30 ng/ml FGF2, and 0.5 mM ascorbic acid (Rossi et al., 2021). EMLOCs were induced in this medium to day 5. At day 7, the EMLOCs were maintained in non-supplemented N2B27. For orbital shaking culture, cells were aggregated and induced at 80 rpm. Speed was reduced to 75 rpm on day 7.

#### EMLOC single-cell dissociation by cold activated protease for scRNAseq

Type here (UB) Day 7 and Day 16 EMLOCs were dissociated on their respective time points in differentiation according to the protocol we previously described (Olmsted and Paluh, 2021a). In brief, ∼25 EMLOCs from each time point were pooled in a 2 ml centrifuge tube and exposed to 1 ml dissociation solution composed of 10 mg/ml Bacillus licheniformis protease and 125 U/ml DNase in ice-cold 1x PBS supplemented with 5 mM calcium chloride. EMLOCs were incubated on ice in dissociation solution and triturated with a P-1000 pipette every 30-60 s for 8 min. Dissociation to single cells was verified by optical inspection and the reaction was terminated by addition of 1 ml ice-cold 1x PBS with 10% fetal bovine serum (FBS). Cells were pelleted by centrifugation at 1,200 x *g* for 5 min, resuspended in fresh 1x PBS/10% FBS, counted, and centrifuged once more. Supernatant was aspirated completely and cells were resuspended in CryoStor CS10 cryopreservation medium to a final concentration of 1×10^6^ cells per ml, filtered through a 40 μm cell strainer, and transferred to a 1.8 ml Nunc cryo-storage tube. Cells were frozen at -80°C overnight and transferred to a liquid nitrogen dewar. When samples from both time points were dissociated and stored, samples were shipped overnight on dry ice to University of Buffalo Genomics and Bioinformatics Core at the New York State Center of Excellence in Bioinformatics and Life Sciences.

#### Single-cell sequencing with CellPlex, cluster annotation and analysis

When samples were received, they were immediately stored at -80°C. On the day of cell capture for sequencing, day 7 and day 16 EMLOC samples were thawed in a 37°C water bath. Individual time point samples were transferred to separate 15 ml tubes. RPMI1640+10% FBS pre-warmed media was added dropwise to a final volume of 10 ml per tube. Cells were centrifuged at 300 *x g* for 5 min. This washing procedure was performed a total of three times. After the final wash, medium was completely removed and cell samples were separately resuspended in 100 μl of Cell Multiplexing Oligo (10x Genomics). The two populations were suspended with two different oligos as directed by the manufacturer’s instructions. After a brief incubation, cells were washed 3x with ice cold 1x PBS (pH 7.4) + 1% bovine serum albumin (BSA). Cells were then resuspended in 250 ul 1x PBS/1% BSA and counted on a Logos Biosystems LUNA II in bright field mode with 0.4% trypan blue. The two cell populations with different barcodes were then pooled to 10,000 cells (5,000 from each time point) and recounted. The combined single cell suspension was provided as input for the 10x Genomics Single Cell v3.1 protocol with Feature Barcode technology. After libraries were prepared, they were loaded onto an Illumina NextSeq in high-output mode with a general target of 50,000 reads per cell to provide for sufficient depth and transcriptomic saturation. Post sequencing, data was demultiplexed and provided as input into the 10x Genomics Cell Ranger multipipeline (ver 4), which quantifies the transcriptomic profile of each cell against a reference genome. Sequence saturation, detected barcodes per cell, percent of transcripts in cell, and general alignment statistics were evaluated for quality. Cell Ranger matrix files were then used as input into the R Bioconductor package Seurat (ver 4). Cells with outlier-status, abnormal gene detection rates, and high mitochondrial transcript load that is an indicator of cellular stress were filtered from the analysis. After filtering, the data was underwent Seurat normalization and integrated using the SCTransform and integration protocol, followed by UMAP (Uniform Manifold Approximation and Projection) or the alternative PHATE dimensionality reduction for visualization (Moon et al., 2019). Using the called clusters, cluster-to-cluster differential expression testing using the Wilcoxon Rank Sum Test was used to identify cluster-defining biomarkers, as well as further exploratory analysis with known biomarker genes (gene lists provided in **Supplementary data file**). Data was then exported from Seurat for further analysis in the Loupe browser.

#### Calcium imaging of contractility

The EMLOCs were incubated with Fluo-4 AM dye as described above in 1 ml of medium for 30 min. Cells were rinsed once in HBSS and imaged in BrainPhys medium without Phenol red. Timelapse series were acquired at 50 ms exposure using a 488 nm LED at 200 ms intervals for 1.5 min duration. Analysis of calcium spike transients was performed using ImageJ. We performed wide field fluorescence microscopy using a Zeiss Axio Observer.Z1 inverted fluorescence microscope (20x/0.8 air objective for live cell calcium imaging). Images were acquired using an Hamamatsu ORCA ER CCD camera and Zeiss AxiovisionRel software (ver. 4.8.2).

#### Phase contrast and whole mount immunofluorescence

Phase contrast microscopy was performed at room temperature directly in the biosafety hood. Images were acquired using a Zeiss Invertoskop 40C (5x/0.12 CP-Apochromat, 10x/0.25 Ph1 A-Plan and 20x/0.30 Ph1 LD A-Plan, 40x/0.50 Ph2 LD A-Plan) mounted with an Olympus DP22 color camera and cellSens acquisition software. Whole-mount immunofluorescence preparation was performed as previously described (Veenvliet et al., 2020; Olmsted and Paluh, 2021). EMLOCs were pooled on the day of fixation, rinsed once with 1x phosphate-buffered saline (PBS), and fixed in 10% neutral buffered formalin solution at 4°C for 2 h. Samples were washed three times in 1x PBS for 5 min at room temperature. Samples were then permeabilized by three successive incubations in 0.2% Triton X-100 in 1x PBS (PBST) for 20 min at 4°C, and blocked overnight in 1% BSA in PBST. For primary antibody incubation, samples were distributed evenly to 12-well plates in 1 ml blocking solution per well. Primary antibodies were added to requisite dilutions in 1% BSA (1x PBS): anti-SOX2 (goat, 5 μg/ml); anti-GATA4 (5 μg/ml); anti-GATA6 (5 μg/ml); anti-CDH1/E-Cadherin (5 μg/ml); anti-FOXA2 (5 μg/ml); anti-β-III-tubulin (rabbit, 1:2,000); phospho-Tau Ser214 (rabbit, 1 μg/ml); CDH2/N-Cadherin (1:200); anti-Collagen Type 1 (1:500, 1 mg/ml stock); anti-Laminin (1:500, 1 mg/ml); anti-Desmin (5 μg/ml); anti-Cardiac Troponin-T (25 μg/ml). Plates were left rocking at 4°C for 24-48 h, rinsed three times in blocking solution, then three times in PBST for 5 min each at room temperature (2 ml centrifuge tubes). Secondary antibodies were incubated 1:1,000 with 2 drops of NucBlue fixed cell stain (Invitrogen) directly in the 2 ml tubes overnight at 4°C. Goat anti-mouse Cy5 secondary antibody was added the next day following washes steps to dilute donkey anti-goat AlexaFluor 594 secondary antibody for samples stained with three antibodies (mouse, rabbit, goat). Samples were again incubated overnight rocking at 4°C. Stained and rinsed EMLO samples were post-fixed in 10% neutral buffered formalin for 20 min at 4°C, and equilibrated in 0.1 M phosphate buffer (PB: 0.025 M NaH_2_PO_4_, 0.075 M Na_2_HPO_4_, pH 7.4) containing 0.2% Triton-X 100 by three successive incubations of 5 min at room temperature. To clear samples, 0.1 M PB was aspirated and replaced with 100 μl of 88% Histodenz solution (w/v) dissolved in 0.2 M PB and filter sterilized. Samples were left in the dark at 4°C for 24 h, mounted on glass slides and sealed in clear nail polish for imaging. Samples were imaged on a Leica confocal TCS SP5 II system in conjunction with Leica Application Suite Advanced Fluorescence software. The SP5 II system was equipped with 10x/0.30 HCX PL FLUOTAR air, 20x/0.70 HC PL APO CS air or immersion, and 40x/1.25 HCX PL APO immersion objective lenses. Complete or partial Z-stacks were acquired at ∼2-2.5 μm separation distance. If necessary, images were corrected linearly for brightness in ImageJ. Maximally projected Z-stacks were performed directly in the Leica software and exported, or were made using Z-project in ImageJ.

### QUANTIFICATION AND STATISTICAL ANALYSIS

#### Quantification of immunofluorescence signal

Quantification immunofluorescence signals along the anterior-posterior axis was performed as described (Rossi et al., 2021). cTnT signal was quantified from maximal projection images (z-axis) and the FOXA2 was quantified from single Z-slices in order to capture the gut tube. The anterior-posterior axis length was measured from pole-to-pole for each gastruloid. Fluorescence intensity was determined using the plot profile tool in Fiji ImageJ, and was normalized along with gastruloid length to enable comparative analysis. FOXA2 and cTnT mean curves were plotted in GraphPad Prism 9 and juxtaposed. Curves were smoothed using a LOWESS function in GraphPad. Only the single channels in question were quantified.

#### Statistical analysis and reproducibility

Microsoft Excel (v16.16.27) and GraphPad Prism 9 (v9.0.2) were used for statistical analysis and data plotting. Data are reported as (mean +/- s.e.m.), analyzed using paired or unpaired two-tailed t-test as indicated. ****p<0.0001, ***p<0.001, **p<0.01, *p<0.05, n.s. not significant (α=0.05 threshold for significance). Power analysis was not performed. Detailed information for each experiment is provided in Results and Figure Legends. Key resources including primary antibodies, chemicals and other reagents, software and equipment, and commercial kits are provided (**Key Resources Table** above).

#### Figures#

Figures for this manuscript were made with a combination of Keynote (v10.3.8), Adobe Illustrator Creative Commons 2020, and BioRender.com. Data were analyzed using Excel and GraphPad Prism 9, and plots were generated using GraphPad Prism 9.

## REFERENCES

Andersen, P., Tampakakis, E., Jimenez, D. V., Kannan, S., Miyamoto, M., Shin, H. K., Saberi, A., Murphy, S., Sulistio, E., Chelko, S. P., et al. (2018). Precardiac organoids form two heart fields via Bmp/Wnt signaling. Nat. Commun. 9, 3140.

Anderson, C., Khan, M.A.F., Wong, F., Solovieva, T., Oliveira, N. M. M., Baldock, R. A. et al. (2016). A strategy to discover new organizers identifies a putative heart organizer. Nat. Commun. 7, 12656.

Ashton, J.L., Burton, R.A.B., Bub, G., Smaill, B.H., Montgomery, J.M. 2018. Synaptic plasticity in cardiac innervation and its potential role in atrial fibrillation. Front. Physiol. 9, 240.

Asp, M., Giacomello, S., Larsson, L., Wu, C., Furth, D., Qian, X. et al. (2019). A spatiotemporal organ-wide gene expression and cell atlas of the developing human heart. Cell 179, 1647–1660.

Beccari, L., Moris, N., Girgin, M., Turner, D.A., Baillie-Johnson, P., Cossy, A-C. et al. 2018. Multi-axial self-organization properties of mouse embryonic stem cells into gastruloids. Nature 562, 272–276.

Campos, I.D., Pinto, V., Souda, N., Pereira, V.H. 2018. A brain within the heart: a review on the intracardiac nervous system. J. Molec. Cell. Cardiol. 119, 1–9.

Chang, E.A., Tomov, M.L., Suhr, S.T., Luo, J., Olmsted, Z.T., Paluh, J.L., Cibelli, J. 2016. Derivation of ethnically diverse human induced pluripotent stem cell lines. Sci. Rep. 5, 15234.

Clevers, H. 2016. Modeling development and disease with organoids. Cell 165, 1586–1597.

Cortes, C., Francou, A., De Bono, C., Kelly, R.G. 2018. Epithelial properties of the second heart field. Circulation Res. 122, 142–154.

Cui, Y., Zheng, Y., Liu, X., Yan, L., Fan, X., Yong, J. et al. Single-cell transcriptome analysis maps the developmental track of the human heart. Cell Rep. 26, 1934–1950.

Das, S., Gordian-Velez, W.J., Ledebur, H.C., Mourkioti, F., Rompolas, P., Chen, H. I. et al. 2020. Innervation: the missing link for biofabricated tissues and organs. NPJ Regen. Med. 5, 11.

Desgrange, A., Le Garrec, J-L., Meilhac, S.M. 2018. Left-right asymmetry in heart development and disease: forming the right loop. Development 145, dev162776.

Drakhlis, L., Biswanath, S., Farr, C-M., Lupanow, V., Teske, J., Ritzenhoff, K. et al. 2021. Human heart-forming organoids recapitulate early heart and foregut development. Nat. Biotechnol. 39, 737–746.

Fedele, L., Brand, T. 2020. The intrinsic cardiac nervous system and its role in cardiac pacemaking and conduction. J. Cardiovasc. Dev. Dis. 7, 54.

Giacomelli, E., Bellin, M., Sala, L., Meer, B.J. van, Tertoolen, L.G.J., Orlova, V.V., Mummery, C.L. 2017. Three-dimensional cardiac microtissues composed of cardiomyocytes and endothelial cells co-differentiated from human pluripotent stem cells. Development 144, 1008–1017

George, R.M., Maldonado-Velez, G., Firulli, A.B. 2020. The heart of the neural crest: cardiac neural crest cells in development and regeneration. Development 147, dev188706.

Han, L., Chaturvedi, P., Kishimoto, K., Koike, H., Nasr, T., Iwasawa, K. et al. 2020. Single cell transcriptomics identifies a signaling network coordinating endoderm and mesoderm diversification during foregut organogenesis. Nat. Commun. 11, 4158.

Harvey, R.P. 2002. Patterning the vertebrate heart. Nat. Rev. Genet. 3, 544–556.

Hasan, W. 2013. Autonomic cardiac innervation: development and adult plasticity. Organogenesis 9, 176–193.

Hofbauer, P., Jahnel, S.M., Papai, N., Giesshammer, M., Deyett, A., Schmidt, C. et al. Cardioids reveal self-organizing principles of human cardiogenesis. Cell 184, 3299–3317.

Hookway, T.A., Matthys, O.B., Mendoza-Camacho, F.N., Rains, S., Sepulveda, J.E., Joy, D.A., McDevitt, T.C. 2019. Phenotypic Variation Between Stromal Cells Differentially Impacts Engineered Cardiac Tissue Function. Tissue Eng. Part A 25, 773–785.

Kidokoro, H., Yonei-Tamura, S., Tamura, K., Schoenwolf, G.C., Saijoh, Y. 2018. The heart tube forms and elongates through dynamic cell rearrangement coordinated with foregut extension. Development 145, dev152488.

Kim, K.H., Nakaoka, Y., Augustin, H.G., Koh, G.Y. 2018. Myocardial angiopoietin-1 controls atrial chamber morphogenesis by spatiotemporal degradation of cardiac jelly. Cell Reports 23, 2455–2466.

Krishnan, A., Samtani, R., Dhanantwari, P., Lee, E., Yamada, S., Shiota, K. et al. 2014. A detailed comparison of mouse and human cardiac development. Pediatr. Res. 76, 500–507.

Lee, J., Sutani, A., Kaneko, R., Takeuchi, J., Sasano, T., Kohda, T. et al. 2020. In vitro generation of functional murine heart organoids via FGF4 and extracellular matrix. Nat. Commun. 11, 4283.

Lind, J.U., Busbee, T.A., Valentine, A.D., Pasqualini, F.S., Yuan, H., Yadid, M. et al. 2017. Instrumented cardiac microphysiological devices via multimaterial three-dimensional printing. Nat. Mater. 16, 303–308.

Ma, Z., Wang, J., Loskill, P., Huebsch, N., Koo, S., Svedlund, F.L. et al. 2015. Self-organizing human cardiac microchambers mediated by geometric confinement. Nat. Commun. 6, 7413.

Macqueen, L.A., Sheehy, S.P., Chantre, C.O., Zimmerman, J.F., Pasqualini, F.S., Liu, X. et al. 2018. A tissue-engineered scale model of the heart ventricle. Nat. Biomed. Eng. 2, 930–941.

Moon, K.R., van Dijk, D., Wang, Z., Gigante, S., Burkhardt, D.B., Chen, W.S. et al. Visualizing structure and transitions in high-dimensional biological data. Nat. Biotechnol. 37, 1482–1492.

Moris, N., Anlas, K., van den Brink, S.C., Alemany, A., Schroder, J., Ghimire, S., et al. 2020. An in vitro model of early anteroposterior organization during human development. Nature 582, 410–415.

Miao, L., Li, J., Lu, Y., Shieh, D., Mazurkiewicz, J.E., Barroso, M. et al. 2019. Cardiomyocyte orientation modulated by the Numb family proteins-N-cadherin axis is essential for ventricular wall morphogenesis. Proc. Natl. Acadm. U S A 116, 15560–15569.

Nascone, N. Mercola, M. 1995. An inductive role for the endoderm in Xenopus cardiogenesis. Development 121, 515–523.

Nguyen, D.C., Hookway, T.A., Wu, Q., Jha, R., Preininger, M.K., Chen, X. et al. 2014. Microscale generation of cardiospheres promotes robust enrichment of cardiomyocytes derived from human pluripotent stem cells. Stem Cell Rep. 3, 260–8.

Olmsted, Z.T., Paluh, J.L. 2021a. Co-development of central and peripheral neurons with trunk mesendoderm in human elongating multi-lineage organized gastruloids. Nat. Commun. 12, 3020. doi: 10.1038/s41467-021-23294-7.

Olmsted, Z.T., Paluh, J.L. 2021b. Generation of human elongating multi-lineage organized (EMLO) gastruloids” Protoc. Exch. DOI: 10.21203/rs.3.pex-1441/v1.

Olmsted, Z.T., Paluh, J.L. 2021c. Stem cell neurodevelopmental solutions for restorative treatments of the human trunk and spine. Front. Cell. Neurosci. 15, 667590.

Polonchuk, L., Chabria, M., Badi, L., Hoflack, J-C., Figtree, G., Davies, M.J., Gentile, C. 2017. Cardiac spheroids as promising in vitro models to study the human heart microenvironment. Sci. Rep. 7, 7005.

Rajendran, P.S., Challis, R.C., Fowlkes, C.C., Hanna, P., Tompkins, J.D., Jordan, M.C.C. et al. 2019. Identification of peripheral neural circuits that regulate heart rate using optogenetic and viral vector strategies. Nat. Commun. 10, 1944.

Rossi, G., Broguiere, N., Miyamoto, M., Boni, A., Guiet, R., Girgin, M. et al. 2021. Capturing cardiogenesis in gastruloids. Cell Stem Cell 28, 230–240.

Sahu, S., Sharan, S. K. 2020. Translating embryogenesis to generate organoids: novel approaches to personalized medicine. iScience 23, 101485.

Sargent, P.B., Garrett, E.N. 1995. The characterization of alpha-bungarotoxin receptors on the surface of parasympathetic neurons in the frog heart. Brain Res. 680, 99–107.

Schultheiss, T.M., Xydas, S., Lassar, A.B. 1995. Induction of avian cardiac myogenesis by anterior endoderm. Development 121, 4203–4214.

Silva, A.C., Matthys, O.B., Joy, D.A., Kauss, M.A., Natarajan, V., Lai, D. et al. 2021. Co-emergence of cardiac and gut tissues promotes cardiomyocyte maturation within human iPSC-derived organoids. Cell Stem Cell 28, 2137–2152.

Steinbeck, J.A., Jaiswal, M.K., Calder, E.L., Kishinevsky, S., Weishaupt, A., Toyka, K.V. et al. 2016. Functional connectivity under optogenetic control allows modeling of human neuromuscular disease. Cell Stem Cell 18, 134–143.

Tom, V.J., Steinmetz, M.P., Miller, J.H., Doller, C.M., Silver, J. 2004. Studies on the development and behavior of the dystrophic growth cone, the hallmark of regeneration failure, in an in vitro model of the glial scar and after spinal cord injury. J. Neurosci. 24, 6531–6539.

Tomov, M.L., Olmsted, Z.T., Dogan, H., Gongorurler, E., Tsompana, M. Out, H.H., et al. 2016. Distinct and shared determinants of cardiomyocyte contractility in multi-lineage competent ethnically diverse iPSCs. Sci. Rep. 6, 37637.

Tyser, R.C.V., Miranda, A.M.A., Chen, C-m., Davidson, S.M., Srinivas, S., Riley, P.R. 2016. Calcium handling precedes cardiac differentiation to initiate the first heartbeat. eLife 5, e17113.

van den Brink, S.C., Baillie-Johnson, P., Balayo, T., Hadjantonakis, A-K., Nowotschin, S., Turner, D.A., Martinez Arias, A. 2014. Symmetry breaking, germ layer specification and axial organization in aggregates of mouse embryonic stem cells. Development 141, 4231–4242.

van den Brink, S.C., Alemany, A., van Batenburg, V., Moris, N., Blotenburg, M., Vivie, J., et al. 2020. Single-cell spatial transcriptomics reveal somitogenesis in gastruloids. Nature 582, 405–409.

van der Linde, D., Konings, E.E.M., Slager, M.A., Witsenburg, M., Helbing, W.A., Takkenberg, J.M., Roos-Hesselink, J.W. e2011. Birth prevalence of congenital heart disease worldwide. J. Amer. Coll. Cardiol. 58, 2241–2247.

Varner, V.D. Taber, L.A. 2012. Not just inductive: a crucial mechanical role for the endoderm during heart tube assembly. Development 139, 1680–1690.

Veenvliet, J.V., Bolondi, A., Kretzmer, H., Haut, L., Scholze-Wittler, M., Schifferl, D. et al. 2020. Mouse embryonic stem cells self-organize intro trunk-like structures with neural tube and somites. Science 370, eaba4937.

