## Supplemental Information for "A Combined Human Gastruloid Model of Cardiogenesis and Neurogenesis"

### Figure S1

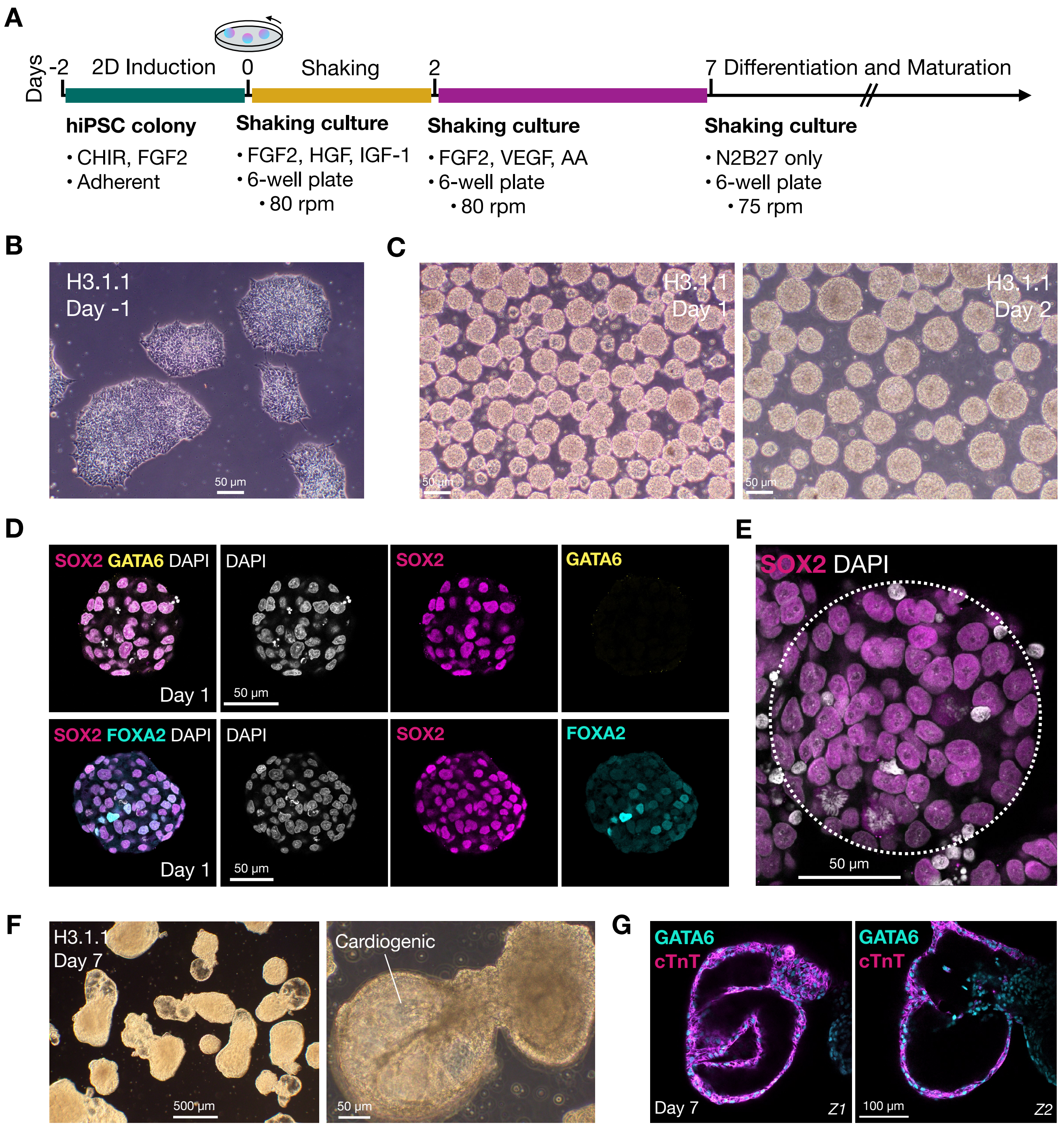

A
Days
-2
2D Induction
0
Shaking
2
7
Differentiation and Maturation
hiPSC colony
• CHIR, FGF2
• Adherent
Shaking culture
• FGF2, HGF, IGF-1
• 6-well plate
• 80 rpm
Shaking culture
• FGF2, VEGF, AA
• 6-well plate
• 80 rpm
Shaking culture
• N2B27 only
• 6-well plate
• 75 rpm
B
H3.1.1
Day -1
50 µm
C
H3.1.1
Day 1
H3.1.1
Day 2
50 µm
50 µm
D
SOX2 GATA6 DAPI
DAPI
SOX2
GATA6
Day 1
50 µm
SOX2 FOXA2 DAPI
DAPI
SOX2
FOXA2
Day 1
50 µm
E
SOX2 DAPI
50 µm
F
H3.1.1
Day 7
500 µm
Cardiogenic
50 µm
G
GATA6
cTnT
Day 7
Z1
GATA6
cTnT
100 µm
Z2

**Figure S1. Directed developmental cardiogenesis and spontaneous contractility in modified neuro-gastro-cardiac EMLOs (EMLOCs), related to Figure 1.** (**A**) Overview of protocol for EMLOC gastruloid generation. Cardiogenesis was induced at 48 h post-aggregation by addition of VEGF and ascorbic acid (AA). (**B**) Phase contrast image of 2D hiPSC colonies 24 h after induction with N2B27 + CHIR/FGF2. (**C**) EMLOC suspension cultures 24 h and 48 h after dissociation and spontaneous aggregation on the orbital shaker. (**D**) Day 1 aggregates do not express GATA6 (yellow) and exhibit non-uniform FOXA2 expression (cyan) by immunofluorescence. (**E**) Immunofluorescence in a single day 1 aggregate (white dotted line) demonstrates uniform, non-polarized expression of SOX2 (magenta). Cells are counterstained with DAPI (grey). (**F**) Phase contrast images of polarized gastruloids with spontaneously contracting cardiogenic chambers (day 7). Annotated EMLOC is shown (right, see also **Movie S2**). (**G**) Two immunofluorescence Z-slices with cardiac biomarkers GATA6 (blue) and cardiac Troponin-T (cTnT, magenta) in a day 7 gastruloid demonstrating chamber cytoarchitecture. Images are representative of the general EMLOC population at these stages. Individual scale bars provided.

### Figure S2

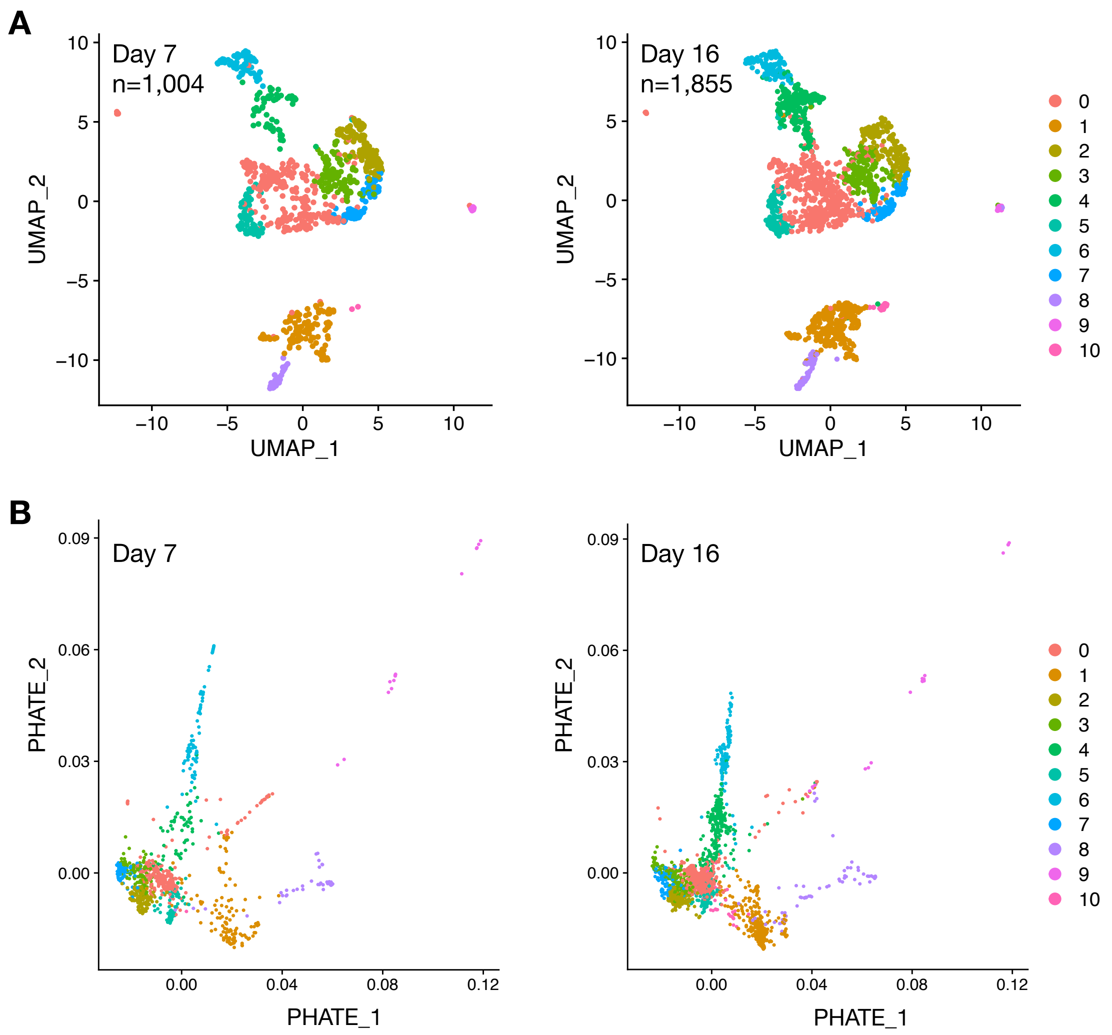
A
Day 7
n=1,004
Day 16
n=1,855
UMAP_1
UMAP_2
B
Day 7
Day 16
PHATE_1
PHATE_2
0
1
2
3
4
5
6
7
8
9
10

**Figure S2. UMAP and PHATE visualization of scRNAseq for day 7 and day 16 EMLOC time points, related to Figure 1.** (**A**) UMAP representation of day 7 (1,004 cells) and day 16 (1,855 cells) according to the integrated dataset with ten clusters. (**B**) PHATE representation of samples in (**A**).

### Figure S3

**A** **Splanchnic mesoderm**

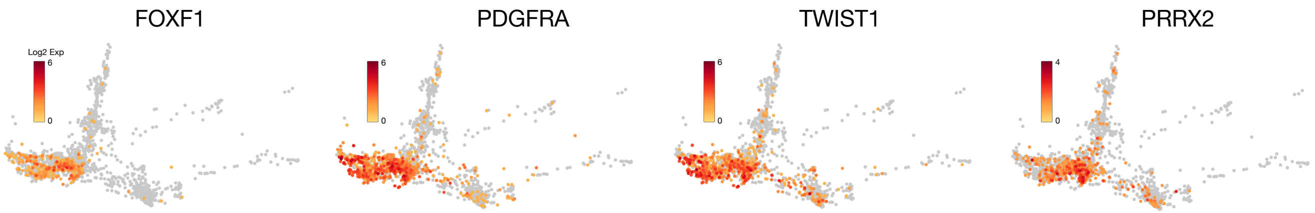


**B** **Ventricular specification and morphogenesis**

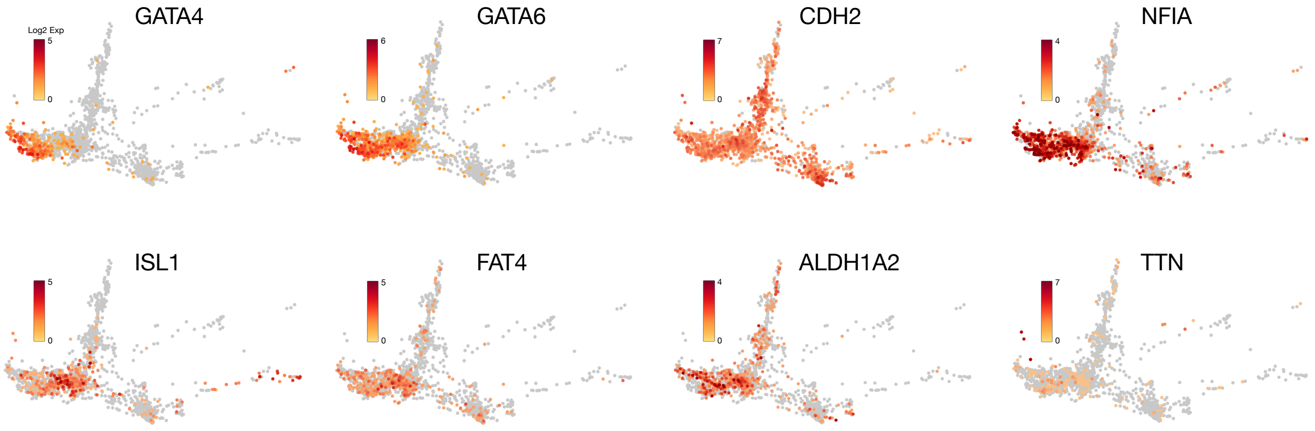


**C** **Cardiac neural crest cells**

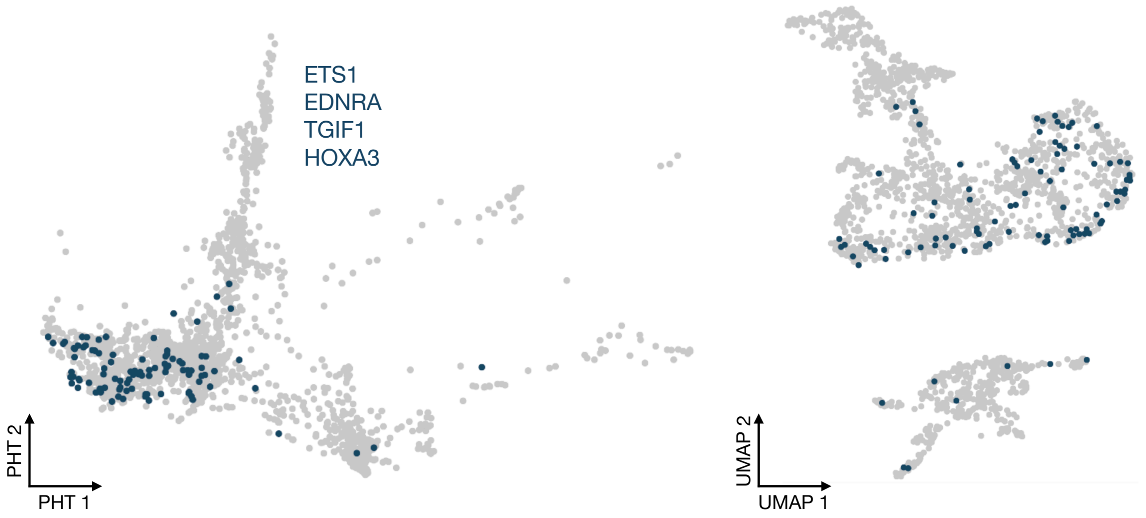

**Figure S3. Developmental features of cardiogenesis in EMLOCs, related to Figure 2.** (**A**) Characteristic gene biomarkers for splanchnic mesoderm (*FOXF1, PDGFRA, TWIST1, PRRX2*). (**B**) Gene biomarkers of ventricular cell specification and morphogenesis (*GATA4, GATA6, CDH2, NFIA, ISL1, FAT, ALDH1A2, TTN*). (**C**) PHATE (left) and UMAP (right) visualization of cells co-expressing *ETS1, EDNRA, TGIF1, HOXA2* consistent with a cardiac neural crest cell phenotype.

#### Figure S4

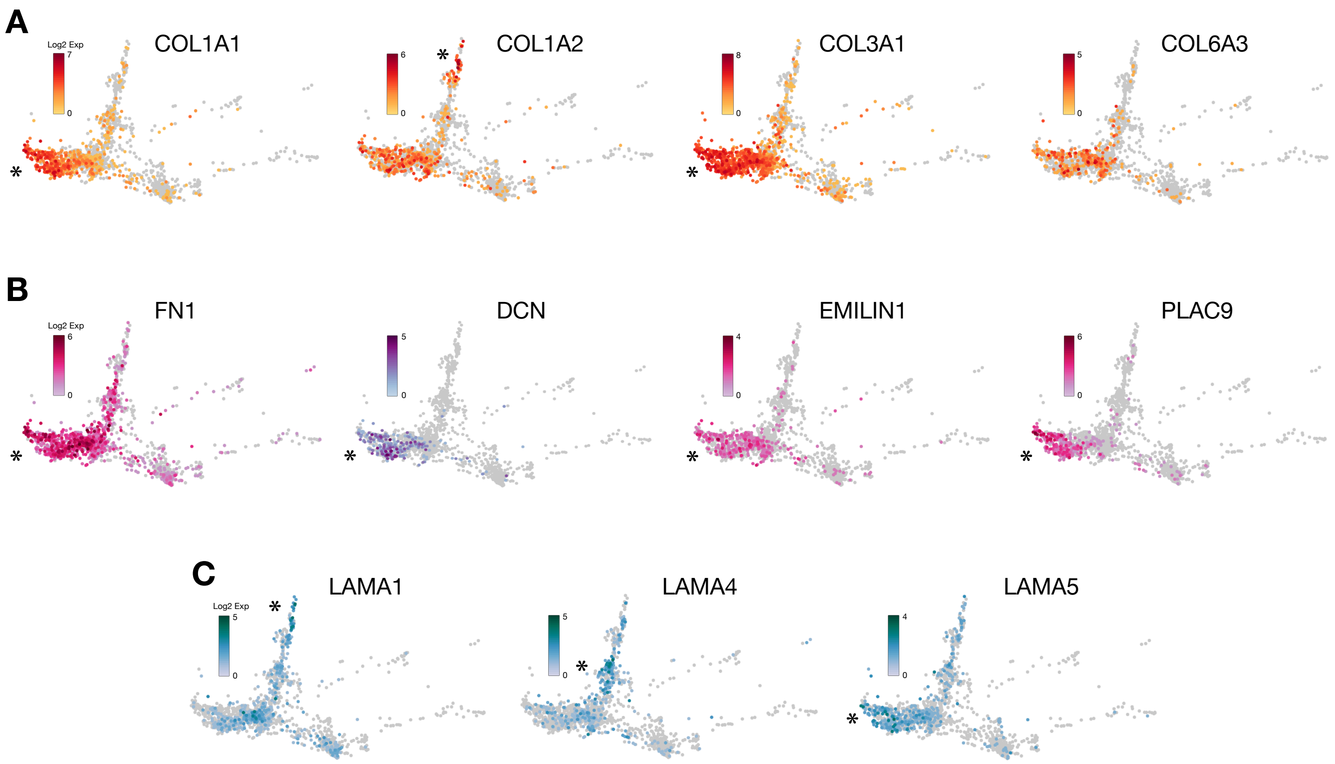
A
COL1A1
Log2 Exp
7
0
*
COL1A2
6
0
*
COL3A1
8
0
*
COL6A3
5
0
B
FN1
Log2 Exp
6
0
*
DCN
5
0
*
EMILIN1
4
0
*
PLAC9
6
0
*
C
LAMA1
Log2 Exp
5
0
*
LAMA4
5
0
*
LAMA5
4
0
*

**Figure S4. EMLOCs generate the appropriate cardiac ECM milieu, related to Figure 2.** (**A**) Four collagen genes (*COL1A1, COL1A2, COL3A1, COL6A3*). (**B**) Glycoprotein (*FN1*) and proteoglycan (*DCN*) genes along with *EMILIN1, PLAC9*. (**C**) Three laminin genes (*LAMA1, LAMA4, LAMA5*) show distinct distributions by PHATE. Asterisk (*) indicates region of highest expression (Cui et al., 2019).

#### Figure S5

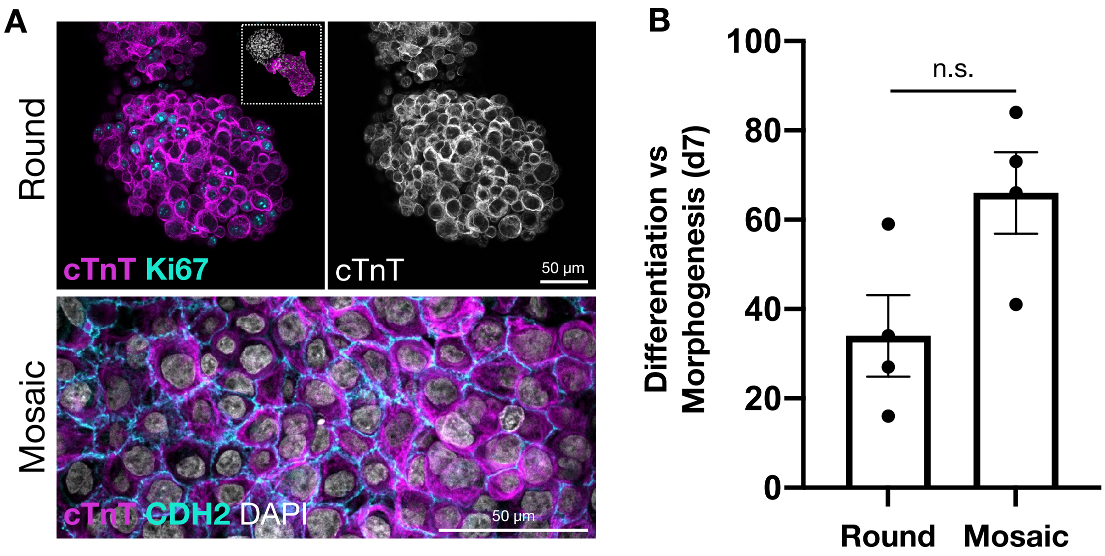
A
Round
Mosaic
cTnT Ki67
cTnT
50 µm
cTnT CDH2 DAPI
50 µm
B
n.s.
Differentiation vs Morphogenesis (d7)
100
80
60
40
20
0
Round
Mosaic

**Figure S5. Assessment of cardiomyocyte morphology with differentiation and morphogenesis states, related to Figure 5.** (**A**) Relative proportions of day 7 EMLOCs with rounded, proliferating cardiomyocyte progenitors versus flattened, mosaic cardiomyocyte progenitors suggests co-existing phases of differentiation and morphogenesis as previously described for mouse cardiogenesis (N = 4 replicate experiments; n.s. p = 0.1776, t = 1.754, df = 3 by paired two-tailed t-test). (**B**) Immunofluorescence examples of rounded, proliferating progenitors (top: cTnT, Ki67) versus flat, mosaic progenitors (bottom: cTnT, CDH2, DAPI). Left inset is example of whole EMLOC with rounded cardiomyocytes. Individual scale bars provided.

#### Figure S6

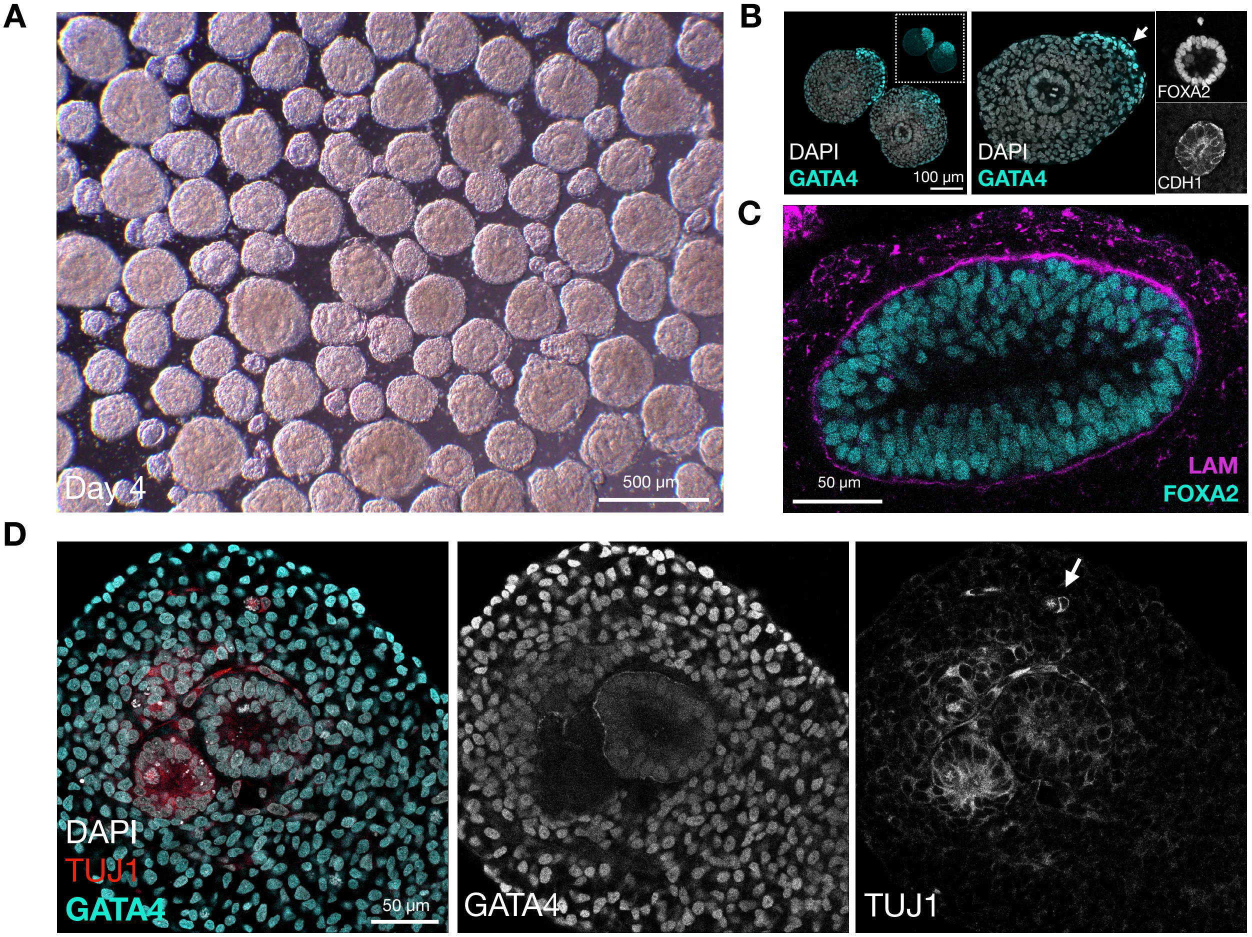

A
Day 4
500 µm
B
DAPI
GATA4
100 µm
DAPI
GATA4
FOXA2
CDH1
C
50 µm
LAM
FOXA2
D
DAPI
TUJ1
GATA4
50 µm
GATA4
TUJ1

**Figure S6. Additional characterization of the gut tube in EMLOCs, related to Figure 5.** (**A**) Day 4 EMLOCs with visible gut tube endoderm. (**B**) FOXA2+/CDH1+ gut tube endoderm is self-organized posterior to the GATA4+ cardiac crescent. White arrow points to early serous lining of the cardiac crescent. (**C**) FOXA2+ gut tube endoderm is laminated, where has TUJ1+/GATA4- neural rosettes are more continuous with surrounding cells, labeled in (**D**). TUJ1 was also observed in mitotic spindle MTs (white arrow). Individual scale bars provided.

**Figure S7**

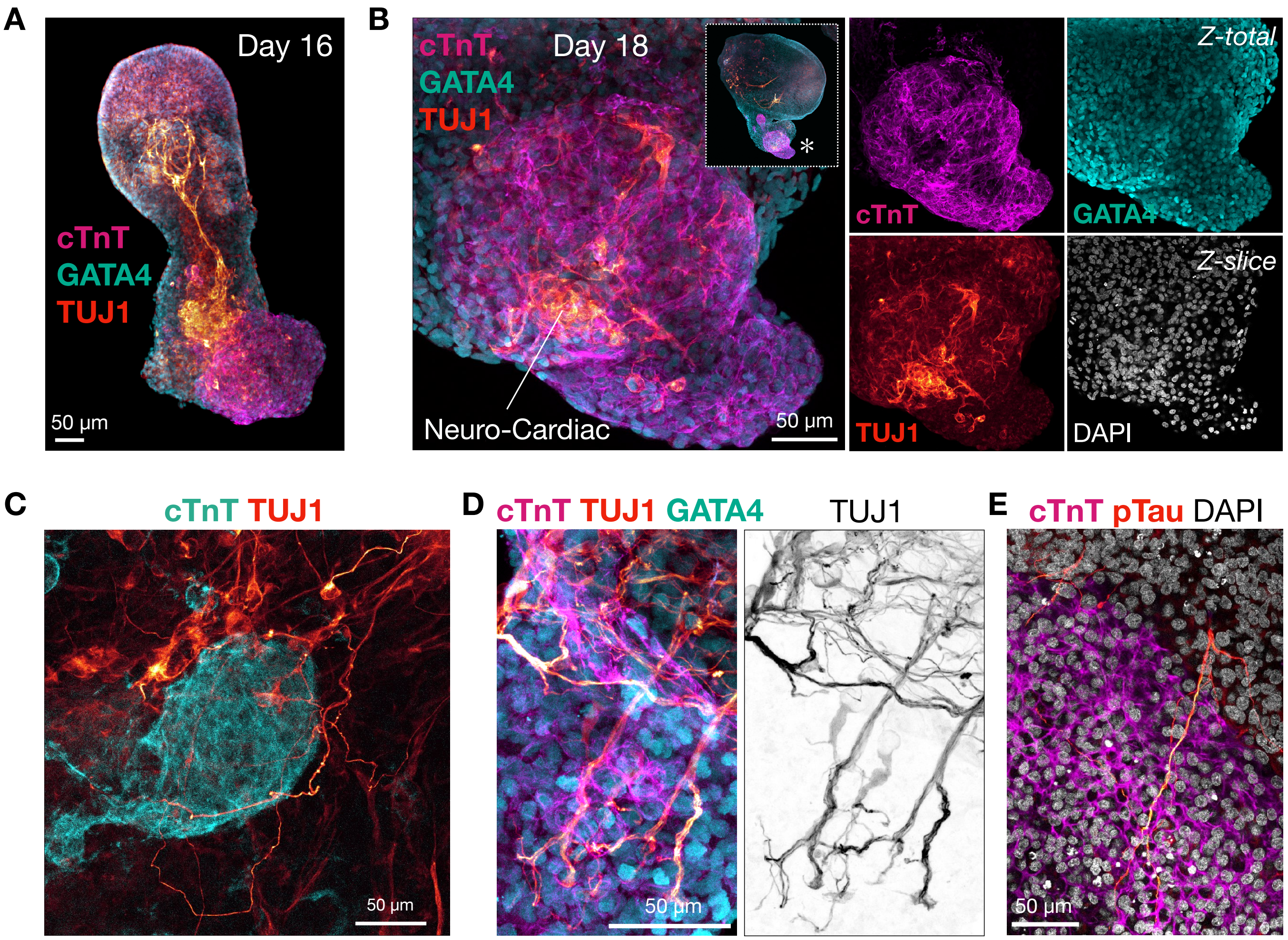
A
Day 16
cTnT
GATA4
TUJ1
50 µm
B
cTnT
GATA4
TUJ1
Day 18
*
Neuro-Cardiac
50 µm
cTnT
GATA4
Z-total
TUJ1
DAPI
Z-slice
C
cTnT TUJ1
50 µm
D
cTnT TUJ1 GATA4
TUJ1
50 µm
E
cTnT pTau DAPI
50 µm

**Figure S7. Additional characterization of neuronal fibers in EMLOCs, related to Figure 8.** (**A**) Immunofluorescence of cardiac biomarkers cTnT (magenta) and GATA4 (cyan) along with TUJ1 (red) in day 16 H3.1.1 EMLOCs demonstrate the emergence of neurons proximal to the cardiac region. (**B**) Immunofluorescence of cTnT (magenta) and TUJ1 (red) at day 18 depicts ganglionated neuronal plexus-like structures in the cardiac region. Merge is shown with individual cTnT, GATA4 (cyan), TUJ1 channels. Left inset is low magnification EMLOC image. (**C**) TUJ1+ neuronal fiber termination (red) onto cTnT+ cells (cyan) reminiscent of nodal innervation. (**D**) Cardiac biomarkers cTnT (magenta) and GATA6 (cyan) with TUJ1. (**E**) Axons identifiable by pTau within the cTnT+ myocardium (right). Individual scale bars provided.

**Table S1. Summary of scRNAseq biomarkers reported in Results, related to Supplementary data file***

| Cell type, tissue type or process | Biomarker reported |
|---|---|
| Cell proliferation | MKI67 |
| First heart field | TBX5 |
| First heart field | NKX2-5 |
| First heart field | HAND1 |
| Second heart field | HAND2 |
| Second heart field | MEF2C |
| Second heart field | TBX18 |
| Second heart field | ISL1 |
| Cardiac sarcomere | TNNT2 |
| Cardiac sarcomere | TNNT1 |
| Cardiac sarcomere | MYL7 |
| Fetal heart development | KRT8 |
| Fetal heart development | KRT18 |
| Fetal heart development | APOE |
| Fetal heart development | PLAC9 |
| Fetal heart development | S100A10 |
| Splanchnic mesoderm | FOXF1 |
| Splanchnic mesoderm | PDGFRA |
| Splanchnic mesoderm | TWIST1 |
| Splanchnic mesoderm | PRRX2 |
| Ventricular specification | GATA4 |
| Ventricular specification | GATA6 |
| Ventricular specification | CDH2 |
| Ventricular specification | NFIA |
| Ventricular specification | ISL1 |
| Ventricular specification | FAT4 |
| Ventricular specification | ALDH1A2 |
| Ventricular specification | TTN |
| Left-right asymmetry | IRX3 |
| Left-right asymmetry | HAND1 |

| | |
|---|---|
| Left-right asymmetry | PITX2 |
| Left-right asymmetry | RTTN |
| Cardiac neural crest cells | ETS1 |
| Cardiac neural crest cells | EDNRA |
| Cardiac neural crest cells | TGIF1 |
| Cardiac neural crest cells | HOXA3 |
| Rapid ventricular conduction | IRX3 |
| Cardiac repolarization | IRX5 |
| Calcium flux | ITPR2 |
| Calcium handling | SLC8A1 |
| AV conduction, organization | GJA1 |
| AV conduction, organization | CACNA1H |
| AV conduction, organization | TBX3 |
| AV conduction, organization | CXCL12 |
| AV conduction, organization | DSP |
| Atrioventricular valves | POSTN |
| Atrioventricular valves | NPR3 |
| Atrioventricular valves | TBX3 |
| Atrioventricular valves | NFATC4 |
| Outflow tract smooth muscle | CNN1 |
| Outflow tract smooth muscle | TAGLN |
| Outflow tract | ISL1 |
| Outflow tract | PDE5A |
| Outflow tract | CDH11 |
| Outflow tract | FN1 |
| Outflow tract | MEGF6 |
| Outflow tract | MSX2 |
| Outflow tract | SEMA3C |
| Outflow tract | EMILIN1 |
| Cardiac jelly | VCAN |
| Cardiac jelly | ADAMTS1 |
| Cardiac jelly | ANGPT1 |
| Vascular endothelial cells | KDR |

| | |
|---|---|
| Vascular endothelial cells | FLT1 |
| Vascular endothelial cells | ESAM |
| Vascular endothelial cells | CDH5 |
| Vascular endothelial cells | GJA4 |
| Epicardial cells | WT1 |
| Epicardial cells | TCF21 |
| Epicardial cells | TPJ1 |
| Epicardial cells | LHX2 |
| Epicardial cells | LHX9 |
| Epicardial cells | TBX18 |
| Epicardial cells | PLAC9 |
| Cardiac fibroblast | IFI16 |
| Cardiac fibroblast | TBX5 |
| Cardiac fibroblast | IGFBP5 |
| Cardiac fibroblast | BTS2 |
| Extracellular matrix | COL1A1 |
| Extracellular matrix | COL1A2 |
| Extracellular matrix | COL3A1 |
| Extracellular matrix | COL6A3 |
| Extracellular matrix | FN1 |
| Extracellular matrix | DCN |
| Extracellular matrix | EMILIN1 |
| Extracellular matrix | PLAC9 |
| Extracellular matrix | LAMA1 |
| Extracellular matrix | LAMA4 |
| Extracellular matrix | LAMA5 |
| Foregut endoderm | FOXA2 |
| Foregut endoderm | NKX2-1 |
| Foregut endoderm | EPCAM |
| Foregut endoderm | CDH1 |
| Foregut endoderm | HHEX |
| Foregut endoderm | SHH |
| Trunk neuroectoderm | SOX2 |

| | |
|---|---|
| Trunk neuroectoderm | FABP7 |
| Trunk neuroectoderm | ZIC1 |
| Trunk neuroectoderm | RFX4 |
| Trunk neuroectoderm | HES5 |
| Trunk neuroectoderm | EDNRB |
| Trunk neuroectoderm | NTRK2 |
| Trunk neuroectoderm | OLIG3 |
| Trunk neuroectoderm | MSX1 |
| Trunk neuroectoderm | HOXC6 |
| Trunk neuroectoderm | HOXC9 |
| Neuronal | STMN2 |
| Neuronal | GAP43 |
| Neuronal | TUBB3 |
| Neuronal | ELAVL3 |
| Neuronal | DLG4 |
| Neuronal | CAMK2A |
| Neuronal | SLC18A3 |
| Neuronal | SLC17A6 |
| Neuronal | CHRNA3 |
| Neuronal | NTRK3 |
| Sensory neuron | POU4F1 |
| Motor neuron | MNX1 |
| Autonomic neuron | PHOX2B |
| Autonomic neuron | ASCL1 |
| Sympathetic neuron | INSM1 |
| Sympathetic neuron | ISL1 |
| Schwann cell glia | SOX10 |
| Schwann cell glia | PLP1 |
| Schwann cell glia | MPZ |
| Schwann cell glia | S100B |
| Schwann cell glia | TFAP2B |
| Schwann cell glia | NGFR |
| Neuro-cardiac patterning | NPY |

| | |
|---|---|
| Neuro-cardiac patterning | BDNF |
| Neuro-cardiac patterning | SEMA3A |
| Neuro-cardiac patterning | PRPH |
| Neuro-cardiac patterning | EDNRA |
| Neuro-cardiac patterning | ISL1 |
| Renal | MAL |
| Renal | SIM1 |
| Renal | EPCAM |
| Renal | TFAP2A |
| Renal | SIX1 |
| Renal | WT1 |
| Renal | ITGA8 |
| Renal | EYA1 |
| Renal | PAX2 |
| Renal | PAX8 |
| Renal | LHX1 |
| Renal | EMX2 |
| BMP signaling | BMP4 |
| BMP signaling | BMP7 |
| WNT signaling | WNT2B |
| WNT signaling | WNT1 |
| WNT signaling | WNT3A |
| Hox code | HOXA4 |
| Hox code | HOXC9 |
| Hox code | HOXD8 |
| Cadherin | CDH1 |
| Cadherin | CDH6 |
| Cadherin | CDH11 |

*An extended version of this table appears as a Supplementary data file
